# Neural adaptation at stimulus onset and speed of neural processing as critical contributors to speech comprehension independent of hearing threshold or age

**DOI:** 10.1101/2023.12.22.573060

**Authors:** Jakob Schirmer, Stephan Wolpert, Konrad Dapper, Moritz Rühle, Jakob Wertz, Marjoleen Wouters, Therese Eldh, Katharina Bader, Wibke Singer, Etienne Gaudrain, Deniz Başkent, Sarah Verhulst, Christoph Braun, Lukas Rüttiger, Matthias H. J. Munk, Ernst Dalhoff, Marlies Knipper

## Abstract

Loss of afferent auditory fiber function (cochlear synaptopathy) has been suggested to occur before a clinically measurable deterioration of subjective hearing threshold. This so-called “hidden” hearing loss is characterized by speech comprehension difficulties. We examined young, middle-aged, and older individuals with and without hearing loss using pure-tone (PT) audiometry, short-pulsed distortion-product otoacoustic emissions (DPOAE), auditory brainstem responses (ABR), auditory steady state responses (ASSR), speech comprehension (OLSA), and syllable discrimination in quiet and noise. After normalizing OLSA thresholds for PT thresholds (“PNOT”), differences in speech comprehension still remained and showed no significant dependence on age, allowing us to categorize participants into groups with good, standard, and poor speech comprehension. Listeners with poor speech comprehension in quiet exhibited smaller firing rate adaptions at stimulus onset (as measured by the difference between DPOAE threshold and pure-tone threshold) and delayed supra-threshold ABR waves I-V, suggesting high spontaneous rate low threshold fiber cochlear synaptopathy. In contrast, when speech comprehension was tested in noise, listeners with poor speech comprehension had larger DPOAEs acceptance rate, putatively resulting from altered basilar membrane compression (recruitment). This was linked with higher uncomfortable loudness levels and larger ASSR amplitudes. Moreover, performance in phoneme discrimination was significantly different below (/o/-/u/) and above the phase-locking limit (/i/-/y/), depending on whether vowels were presented in quiet or ipsilateral noise. This suggests that neural firing rate adaptation at stimulus onset is critical for speech comprehension, independent of hearing threshold and age, whereas the recruitment phenomenon counterbalances the loss in speech-in-noise discrimination due to impaired threshold.

**Significance Statement:** Age-related hearing loss is the third largest modifiable risk factor for cognitive decline. It has been suggested that the link between hearing loss and cognitive decline is not fully explained by hearing threshold loss. We here suggest that language comprehension deficits may be used as an early indication of future hearing loss and therefore cognitive decline. We found that, independent of age and pure-tone thresholds, speech comprehension in quiet and ipsilateral noise depend on different onset firing-rate adaptations of inner hair cells (measured by DPOAE threshold), along with cochlear synaptopathy of high spontaneous rate auditory nerve fibers and neural spiking synchronicity. These measures may be used as possible future indicators of risk for cognitive decline.

## INTRODUCTION

Age-related hearing loss is the most prevalent disorder of aging and is associated with future cognitive impairment (Livingston et al., 2017). Recent studies indicate that the association between hearing and cognition also exists in individuals with subclinical hearing loss—that is, those with normal pure-tone audiograms < 25 dB (Golub et al., 2020; Hoppe et al., 2022). This suggests that the reason for the cognitive decline after hearing loss is not necessarily due to a loss of hearing threshold. Here, we suggest that afferent auditory fiber loss (cochlear synaptopathy) is a candidate contributor, as it may actually precede loss of outer hair cells **(OHCs)** and an overt threshold loss, as shown in animals (Sergeyenko et al., 2013; Mohrle et al., 2016; Monaghan et al., 2020) and predicted for humans (Plack et al., 2014; Viana et al., 2015; Kobel et al., 2017; Liberman and Kujawa, 2017; Bharadwaj et al., 2019; Mepani et al., 2021), (Frisina, 2009; Fullgrabe et al., 2014; Bharadwaj et al., 2019). Specifically, the damage of low spontaneous rate **(SR)** high threshold auditory nerve fibers **(ANFs)** is suggested to be responsible for cochlear synaptopathy linked to speech comprehension deficits, as these ANFs play a critical role in coding supra-threshold sound features as speech in noise (Plack et al., 2014; Chambers et al., 2016; Hesse et al., 2016; Liberman and Kujawa, 2017; Asokan et al., 2018; Bakay et al., 2018; Wu et al., 2019; Monaghan et al., 2020). Broadband sounds like speech are decomposed into narrowband signals that vary depending on if nerve spikes are phase-locked to the envelope (above phase locking limit **(PLL)**) or phase-locked to the stimulus (below PLL) (Moore, 2021), possibly due to differential contributions of the two ANF types (Huet et al., 2018). Frequency band specific deficits independent of threshold should therefore be regarded in the context of cochlear synaptopathy. Moreover, in patients with normal audiograms, speech comprehension deficits have been linked to deficits in extended high frequency (> 10 kHz; **EHF**) hearing (Apoux and Bacon, 2004; Le Prell et al., 2013; Levy et al., 2015; Vitela et al., 2015; Moore et al., 2017; Motlagh Zadeh et al., 2019; Hunter et al., 2020; Song et al., 2022) and central desynchronization (Pichora-Fuller et al., 2007; Frisina, 2009; Bajin et al., 2022).

We measured pure-tone thresholds **(PTTs)** in a total of 89 young, middle-aged, and older individuals. Speech comprehension was tested via speech reception thresholds **(SRT)** in quiet or at a fixed noise level using the standard German Matrix test Oldenburger Satztest **(OLSA)** for either unfiltered “broadband” speech **(OLSA-BB)**, low-pass filtered speech **(OLSA-LP)**, or high-pass filtered speech **(OLSA-HP)**, as described (Garrett et al., 2020). Using a multivariate regression model based on principal component analysis **(PCA)**, participants with matched PTTs were classified into groups with good, standard, and poor speech comprehension. Temporal precision of auditory coding (auditory steady state response, **ASSR)**, estimated distortion-product thresholds (Zelle et al., 2017) that appeared to be a metric for onset firing-rate adaptations of inner hair cells if compared to pure-tone threshold, auditory brainstem responses (**ABR** wave I-VI amplitude and latency), and discrimination of vowel contrasts below or above PLL were analyzed. We strikingly found that onset firing rate adaptation, high-SR ANF cochlear synaptopathy, and recruitment influences on neural spiking synchronicity contribute to speech comprehension independent of age and threshold.

## METHODS

The study was conducted at the Department of Otolaryngology of the University of Tübingen and approved by the ethics committee of Tübingen University (faculty of medicine) (ethical approval-number 392/2021BO2). Written informed consent was given by all participants. All methods were used according to the Declaration of Helsinki’ by the World Medical Association **(WMA)** for human research ethics.

### Participants

We recruited 112 participants aged between 18 and 76 years. A checklist to inquire for any of comorbidities described in the ethics protocol was used for study exclusion. Among these were: other hearing-related conditions such as tinnitus, or previous ear-surgery, as well as systemic diseases known to affect hearing. To the end, 89 participants were included in the subsequent analysis, while the remaining ones were excluded due to comorbidities (threshold elevation beyond 40 dB in one or more frequencies and tinnitus) or because of lacking compliance. The included 89 participants were evenly distributed across 3 age-groups, young (18-29 years, n=29), middle-aged (30-55 years, n=32) and older (56-76 years, n=28) (Tab. 1). Participants’ age, gender, handedness and confirmation of normal middle ear function by tympanometry are provided in (Tab. 1). Of the 89 participants, only 63 could be measured for all noise conditions of the German word matrix test OLSA such that in addition to “in quiet” two noise conditions as ipsi- and contralateral noise were obtained, the remaining 26 participants were only tested with the quiet condition.

**Table 1.** Proband information to age, gender (female (f), handedness and results from depression evaluation according to Beck Depression Inventory II (BDI-II), Geriatric Depression Scale (GDS) as well as dementia test Mini Mental Status Test (MMST) and Tympanometry results. Age(years), gender (female=f, male=m), handedness (R=right; L=left). Psychometric results from depression evaluation according to Beck’s Depression Inventory II (BDI-II), Geriatric Depression Scale (GDS) as well as dementia test Mini Mental Status Test (MMST). Tympanometry results describing “Type A” = normal; “Type B” = flat; “Type C” = compliance peak indicating negative pressure; “Type AD” = compliance peak elevated and broader than normal; “Type AS” = reduced compliance peak with normal pressure.

| Proband | Age | Gender | Handedness | BDI* | GDS* | MMST | Tympanometry |
| --- | --- | --- | --- | --- | --- | --- | --- |
| CS001 | 26 | f | right | minimal | minimal | 30 | Type A |
| CS002 | 63 | f | right | minimal | minimal | 30 | Type A |
| CS003 | 59 | m | right | minimal | minimal | 30 | Type A |
| CS004 | 62 | m | right | minimal | minimal | 30 | Type A |
| CS005 | 55 | f | right | minimal | minimal | 30 | Type A |
| CS006 | 60 | f | right | mild | minimal | 30 | Type A |
| CS007 | 38 | f | right | minimal | minimal | 30 | Type A |
| CS008 | 48 | f | right | moderate | mild/moderate | 30 | Type B |
| CS009 | 21 | f | right | minimal | minimal | 30 | Type A |
| CS010 | 47 | f | right | minimal | minimal | 30 | Type A |
| CS011 | 19 | f | right | minimal | minimal | 30 | Type C |
| CS012 | 22 | m | right | minimal | minimal | 29 | Type A |
| CS013 | 26 | f | right | minimal | minimal | 30 | Type A |
| CS014 | 25 | m | right | minimal | minimal | 30 | Type A |
| CS015 | 25 | m | right | mild | minimal | 30 | Type A |
| CS018 | 24 | m | right | minimal | minimal | 29 | Type A |
| CS023 | 69 | m | right | moderate | mild/moderate | 29 | Type A |
| CS025 | 64 | f | right | minimal | minimal | 28 | Type A |
| CS027 | 60 | f | right | minimal | minimal | 30 | Type A |
| CS029 | 64 | m | right | minimal | minimal | 29 | Type A |
| CS030 | 23 | f | right | minimal | minimal | 30 | Type A |
| CS031 | 53 | m | right | minimal | minimal | 29 | Type A |
| CS033 | 26 | f | right | minimal | minimal | 29 | Type A |
| CS034 | 24 | f | right | minimal | minimal | 29 | Type A |
| CS035 | 45 | m | right | minimal | minimal | 28 | Type A |
| CS037 | 45 | m | right | minimal | minimal | 29 | Type A |
| CS038 | 57 | f | right | minimal | minimal | 30 | Type A |
| CS039 | 18 | f | right | minimal | minimal | 29 | Type A |
| CS040 | 57 | f | right | minimal | minimal | 30 | Type A |
| CS041 | 58 | f | right | minimal | minimal | 30 | Type A |
| CS042 | 54 | f | right | mild | minimal | 29 | Type AS |
| CS044 | 62 | f | right | minimal | minimal | 30 | Type AS |
| CS047 | 55 | m | right | minimal | minimal | 29 | Type A |
| CS050 | 62 | f | right | minimal | minimal | 29 | Type A |
| CS051 | 29 | m | left | mild | minimal | 30 | Type AS |
| CS053 | 19 | m | right | moderate | mild/moderate | 30 | Type A |
| CS056 | 28 | f | right | minimal | minimal | 30 | Type A |
| CS057 | 28 | f | right | minimal | minimal | 29 | Type AD |
| CS058 | 21 | f | right | minimal | minimal | 30 | Type A |
| CS059 | 29 | m | left | minimal | minimal | 29 | Type A |
| CS060 | 24 | f | right | minimal | minimal | 29 | Type AD |
| CS061 | 58 | f | right | minimal | minimal | 30 | Type A |
| CS062 | 52 | f | right | minimal | minimal | 30 | Type A |
| CS063 | 28 | m | right | minimal | minimal | 30 | Type A |
| CS065 | 23 | f | right | minimal | minimal | 29 | Type A |
| CS066 | 70 | m | right | minimal | minimal | 30 | Type AD |
| CS067 | 33 | m | right | mild | minimal | 30 | Type AD |
| CS068 | 52 | f | right | minimal | minimal | 30 | Type A |
| CS069 | 24 | f | right | minimal | minimal | 30 | Type A |
| CS070 | 20 | f | left | mild | mild/moderate | 28 | Type A |
| CS071 | 60 | f | right | minimal | minimal | 30 | Type A |
| CS073 | 43 | f | right | minimal | minimal | 29 | Type A |
| CS074 | 21 | m | right | moderate | mild/moderate | 29 | Type A |
| CS075 | 54 | f | right | minimal | minimal | 30 | Type A |
| CS076 | 30 | m | right | minimal | minimal | 30 | Type A |
| CS077 | 52 | f | right | minimal | minimal | 30 | Type A |
| CS078 | 35 | m | right | minimal | minimal | 30 | Type A |
| CS079 | 34 | f | right | minimal | minimal | 28 | Type A |
| CS080 | 35 | m | left | minimal | minimal | 30 | Type A |
| CS081 | 54 | f | right | minimal | minimal | 30 | Type A |
| CS082 | 48 | f | right | minimal | minimal | 29 | Type AD |
| CS083 | 76 | m | right | minimal | minimal | 29 | Type A |
| CS084 | 54 | f | right | minimal | minimal | 30 | Type A |
| CS085 | 40 | f | right | minimal | minimal | 28 | Type A |
| CS086 | 56 | f | right | minimal | minimal | 30 | Type AD |
| CS087 | 32 | f | right | minimal | minimal | 29 | Type A |
| CS088 | 30 | f | right | minimal | minimal | 29 | Type A |
| CS089 | 53 | f | right | minimal | minimal | 30 | Type A |
| CS090 | 56 | f | right | minimal | minimal | 30 | Type A |
| CS091 | 30 | f | right | minimal | minimal | 30 | Type A |
| CS093 | 40 | m | right | minimal | minimal | 30 | Type A |
| CS095 | 50 | m | right | minimal | minimal | 29 | Type A |
| CS096 | 51 | f | right | minimal | minimal | 30 | Type AS |
| CS097 | 40 | f | right | minimal | minimal | 30 | Type A |
| CS098 | 54 | f | right | minimal | minimal | 30 | Type A |
| CS099 | 27 | f | right | minimal | minimal | 29 | Type A |
| CS100 | 25 | m | right | minimal | minimal | 30 | Type A |
| CS101 | 46 | f | right | minimal | minimal | 30 | Type A |
| CS102 | 21 | m | right | mild | minimal | 30 | Type A |
| CS103 | 28 | f | right | minimal | minimal | 28 | Type A |
| CS104 | 68 | f | right | minimal | minimal | 30 | Type AS |
| CS105 | 65 | m | left | minimal | minimal | 30 | Type A |
| CS106 | 23 | f | left | minimal | minimal | 30 | Type A |
| CS107 | 64 | m | right | minimal | minimal | 30 | Type B |
| CS108 | 70 | f | right | minimal | minimal | 29 | Type A |
| CS109 | 70 | w | right | minimal | minimal | 29 | Type A |
| CS110 | 72 | m | right | minimal | minimal | 29 | Type AS |
| CS111 | 62 | m | right | minimal | minimal | 30 | Type A |
| CS112 | 71 | f | right | minimal | minimal | 29 | Type As |

### Neuropsychiatric Scores

To evaluate neuropsychiatric factors as population characteristics which can interfere with sensory functions like hearing, we screened for depression and dementia. For detecting depression, we applied two validated questionnaires: “Becks Depression Inventar II” **(BDI)**, shown to screen for depression in a clinical setting (Beck et al., 1996) and the Geriatric Depression Scale **(GDS)** with focus on affective and cognitive domains (Yesavage et al., 1982). For excluding cognitive decline, we used the gold standard for minimal neuropsychological testing in form of a German version of the Mini-Mental State Examination **(MMSE)** (Folstein et al., 1975; Tombaugh and McIntyre, 1992) to screen and exclude dementia. Within this test the participants are asked for items like orientation in space and time, word short-term memory, subtracting, attentive listening, spelling, reading, writing, executive tests, visuo-construction. With a custom questionnaire we evaluated hearing ability with a self-assessment of hearing in various conversation settings and general information like education and handedness (Maele et al., 2021).

Overall, the analysis of the results of BDI, GDS, and MMSE across age shows that older participants did not score higher than younger subjects. In contrast, younger participants were more affected by moderate depression than older subjects (Fig.1, Tab. 2). Also, no MMSE scored below 27, an indication of intact cognitive function of all participants. Therefore, regarding both depression scales, up to moderate depression was present in four to five subjects from our sample, while cognitive decline and dementia could be excluded (Fig. 1, Tab. 2).

**Figure 1.**
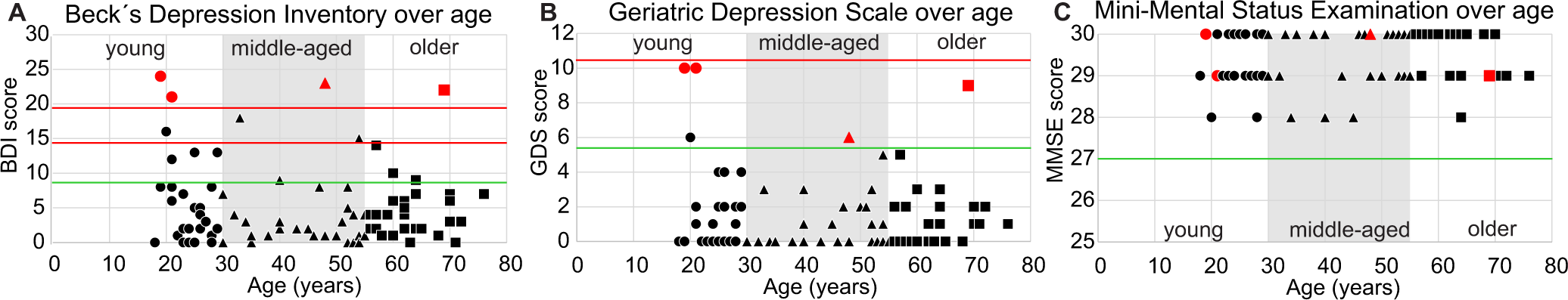
Psychometric results for screening depression (BDI II / GDS) and cognitive decline (MMSE). The ordinates represent the scores and the abscissae the age of the participants. The grey shade indicates the middle-aged group; data points of each group are also represented by different symbols. Colored lines reflect limits for suspecting depression or cognitive decline: green lines mark the borderline for normal scores, red lines the borderline between mild-to-borderline and moderate degree of depression, for the GDI differentiating between mild-to-moderate and severe depression. Note that for MMSE the scaling is reversed with respect to the depression scales: numbers below 27 indicate a pathologic state.

**Table 2.** Psychometric scores for depression and dementia screening. Mean, median and SEM are given for the results of BDI II, GDS and MMSE for all three age groups.

|  | BDI |  |  | GDS |  |  | MMSE |  |  |
| --- | --- | --- | --- | --- | --- | --- | --- | --- | --- |
|  | mean | median | SEM | mean | median | SEM | mean | median | SEM |
| Young | 6.06 | 4 | 1.18 | 1.96 | 1 | 0.47 | 29.46 | 30 | 0.11 |
| Middle | 3.96 | 2 | 1.01 | 0.93 | 0 | 0.27 | 29.56 | 30 | 0.12 |
| Older | 4.75 | 4 | 0.84 | 1.41 | 1 | 0.34 | 29.62 | 30 | 0.10 |

### Otoscopy and impedance audiometry

The ear examination was carried out by ENT physicians from the Department of Otolaryngology, Head and Neck Surgery at the University of Tübingen. Tympanometry and stapedial-reflex measurements were performed with an AT235 (Interacoustics, Middelfart, Denmark) tympanometry system using a 226 Hz stimulus to ensure intact middle-ear transmission (Shanks, 1984) and generally intact neural pathways (Margolis and Goycoolea, 1993).

### Pure-tone audiometry

Pure-tone thresholds (PTT) were measured for air and bone conduction, as well as the uncomfortable loudness level (ULL) using the AT 1000 Audiometer (Auritec, medizindiagnostische Geräte Gmbh, Hamburg, Germany). Bone conduction at 0.25, 0.5, 1, 1.5, 2, 4, 6 kHz was measured using a B71Wbone transducer (Radioear, Middelfart, Denmark). The default pure tone audiometric thresholds from 0.125 to 10 kHz and the ULL (0.25, 0.5, 1, 2, 4, 6 kHz) were measured with Beyerdynamic AT1350A on-ear headphones (Beyerdynamic, Heilbronn, Germany). In addition, extended high frequencies (EHF) thresholds were measured with Sennheiser HDA300 (Sennheiser, Wedemark-Wennebostel, Germany) on-ear headphones at frequencies 11.2, 12.5, 14, 16 kHz. All measurements were conducted in a sound-proof chamber (Industrial Acoustics Company GmBH, Niederkrüchten, Germany).

Pure-tone averages of low frequencies (**PTA**-LF; 0.125, 0.25, 0.5, and 1 kHz), high frequencies (**PTA-HF**; 6, 8, and 10 kHz), extended high frequencies (**PTA-EHF**; 11.3, 12.5, 14, and 16 kHz) and **PTA4** (0.5, 1, 2, and 4 kHz) were derived from the right ear thresholds. These specific PTA groups were chosen to analyze thresholds for specific frequency ranges: (i) PTA-LF, for hearing below the phase-locking limit (PLL) i. e. < 1500 Hz in humans, (ii) PTA4, because this is the worldwide standard for defining hearing-loss severity, (iii) PTA-HF, because these frequencies are regularly measured in the clinical setting, but are above the frequency range of the PTA4, and (iv) PTA-EHF, because those frequencies are suggested to impact speech perception as well (Levy et al., 2015; Hunter et al., 2020), and are above what is usually measured in a clinical setting.

### Auditory Brainstem Responses

The ABR measurements were performed with two-channel recordings using three electrodes (Neuroline 720, Ambu, Bad Nauheim, Germany) with electrode impedance consistently below two kΩ (ground: Fpz - above the nasion; reference - inverting input (-): Fz - hairline; non-inverting input (+): mastoid). As an amplifier, the actiCHamp Plus64 (Brain Products GmbH, Gilching, Germany) was set up according to manufacturer specifications and at a sampling rate of 50 kHz.

Acoustic click stimuli (83µs) were presented at two different stimulus levels (70 dB SPL and 80 dB SPL) with 3000 repetitions of alternating polarity. Stimuli were generated with a Scarlet Focusrite 8i8 gen 3 (Focusrite, United Kingdom) sound-card and presented through ER2 transducers and disposable ER1-14A earpieces (Etymotic Research, Elk Grove Village, Illinois, USA). The participants lay on their backs during the measurements to minimize muscle effects.

After band-pass filtering between 30-2000 Hz (first order FIR filter, Hamming windowed), ABR waveform components were averaged at both stimulus levels. Wave V was determined to be the most prominent peak, typically appearing at 5-6 ms after stimulus onset. Waves I, II, III and VI were then assigned to peaks at 1-2 ms, 2-3 ms, 3-4 and 6-7 ms after stimulus onset respectively. Wave amplitudes were calculated in µV as the difference between leading positive and trailing negative deflections/peaks as described (Hofmeier et al., 2018; Hofmeier et al., 2021). Their latency was determined from the leading positive peak.

### Auditory Steady State Response

After measuring the Auditory Brainstem Response (ABR), the Auditory Steady State Response (ASSR) was measured using the same recording setup without changing the position of the participants. The modulation frequency was set to 116 Hz (rectangular 100% amplitude modulation as described in (Vasilkov et al., 2021) and two blocks of 800 epochs each were recorded at carrier frequencies of 4 and 6 kHz at 70 dB SPL rms. The stimulus duration was set to 400 ms with an epoch duration of 500±10 ms. Responses from all epochs were averaged and the spectral power calculated by FFT (Matlab 2021b). ASSR peak amplitudes (µV) were averaged for the first three harmonics (Vasilkov et al., 2021). Measurements with inadequate signal-to-noise ratios (SNR < 2) or ASSR peak amplitudes > 0.15µV, were excluded from the statistical evaluation.

### Distortion-Product Otoacoustic Emissions

Short-pulsed distortion-product otoacoustic emissions **(p-DPOAE)** were measured to characterize the pre-neural state of the cochlea. Using a pulsed waveform for the second primary **(f_2_)**, along with onset decomposition (Zelle et al., 2017), a technique to capture the short-latency nonlinear-distortion **(ND)** component of the DPOAE (Shera and Guinan, 1999), artefactual interference effects with the longer latency component can be safely avoided (Zelle et al., 2016). p-DPOAE input-output **(I/O)** functions were measured at 8 frequencies (f_2_=0.8, 1.2, 1.5, 2, 3, 4, 6, 8 kHz) using an adaptive algorithm comprising at least four p-DPOAE values. Estimated distortion-product thresholds (L_EDPT_) were derived from these ND-DPOAE based on linear regression to the semi-logarithmically scaled ND-DPOAE-I/O functions (p_DP_ [µPa] vs. L_2_ [dB SPL], cf. (Boege and Janssen, 2002; Zelle et al., 2017). The L_EDPT_ derived from ND-DPOAE I/O functions as well as so-called p-DPOAE level maps have been shown to correlate in an almost 1:1 relationship with auditory thresholds with a hitherto exceptionally small standard deviation of the residual of approximately 6 dB (Zelle et al., 2017; Zelle et al., 2020) over a frequency range of at least 1–8 kHz, and to be highly reproducible (Bader et al., 2021).

p-DPOAE were measured unilaterally with ER-10C probes (Etymotic Res., Elk Grove Village, IL). Calibration of the speakers and the microphone included correction for probe-to-tympanic membrane transfer function based on an artificial ear simulator (B&K type 4157, Bruel & Kjær, Nærum, Denmark) and thus are considered to correspond to sound pressure at the tympanic membrane; for details of calibration procedures, see (Zelle et al., 2017). In order to be accepted, the linear regression to the experimentally obtained I/O function must fulfill several criteria, the most important being related to the quality of the regression procedure, namely regarding the squared correlation coefficient, r^2^ ≥ 0.8, and the standard deviation of the L, σ ≤ 10 dB (for further details, see (Zelle et al., 2017). The acceptance rate reported in the Results Section is then the ratio between accepted I/O functions within a group of n_g_ members across all eight frequencies and both ears, i.e. n_acc_/(2*8*n_g_). Acceptance rate depends on DPOAE amplitudes, background noise, and the linearity of the measured I/O functions, to mention some important contributors. Two sets of interleaved presentations of four two-tone pulse pairs each were averaged over 100 ensembles, where one ensemble consists of 4 blocks with suitable phase shifts yielding a time-domain DPOAE response with vanishing contribution of the two stimulus tones, when averaged; this method is called primary-tone phase variation technique (Whitehead et al., 1996). The first set comprised presentations with f_2_=0.8, 1.5, 3, and 6 kHz in a block of 180 ms length, the second with f_2_=1.2, 2, 4, and 8 kHz in a block of 120 ms length. The adaptive procedure started with L_2_=45 dB SPL for each f_2_-pulse, followed by lower or higher stimulus levels depending on SNR, a minimum L_2_ step size of 3 dB, and estimations based on population data of I/O slopes (Krokenberger, 2019). To be accepted for the subsequent DPOAE threshold estimation, a single ND-DPOAE value requires signal to noise ratios **(SNR)** ≥ 10 dB. L_1_ values are chosen according to a frequency-specific scissor paradigm (Zelle et al., 2017) aimed to equalize both travelling-wave amplitudes of the stimulus tones f_1_ and f_2_ at the f_2_-place in the cochlea (cf. (Zelle et al., 2020)). f_1_-pulses were switched on 2.5 ms after corresponding f_2_-pulses and presented with widths between 40 to 10 ms adjusted to ensure that intracochlearly the correspondent travelling wave has reached a steady state before the f_2_-pulse is switched on. f_2_-pulses had half-maximum widths decreasing from 11.9 to 3.0 ms with ramps decreasing from 4.6 to 1.2 ms for frequencies between 0.8 and 8 kHz, designed to roughly allow settling of the ND component of the DPOAE before the second component of the DPOAE, the coherent-reflection component, builds up, so that both components can be visually distinguished in the time domain (Zelle et al., 2013).

### Speech reception thresholds (OLSA)

Speech intelligibility was tested using the “Oldenburger Satz Test” (OLSA), the German version of the International Matrix test (Wagener et al., 1999; Brand and Kollmeier, 2002), applying three different noise and three different filtering conditions. For each noise condition, participants were presented with 20 sentences each, i.e. in unfiltered broadband noise (OLSA-BB), in low-pass filtered noise (OLSA-LP; frequency components >1500 Hz deleted from the OLSA power spectrum) or high-pass filtered speech (OLSA-HP, <1500 Hz deleted from the OLSA power spectrum; see Fig. 2) (see for details, (Garrett et al., 2020) with the order of the 9 conditions presented in random order. Sentences consisted of 5 words, with a name, verb, number, adjective, and object, and each keyword having 10 response possibilities, producing a large number of stimuli (10^5) combinations from an inventory of a total of 50 words. The speech material of the OLSA is spoken by a male speaker (Wagener et al., 1999) the average F0 of which we determined to be 116 Hz. The target sentences and the masker noise were presented monoaurally (speaker always right, noise same or different ears) over ER2 transducers (see Methods, Auditory Brainstem Responses). The level of the target sentence varied and was decreased after a correct response (i.e., increasing difficulty) or increased after an incorrect response (i.e., decreasing difficulty). The masker noise has been derived from the speech material by randomly shifted overlapping and thus exhibits the same long-term spectrum (Wagener et al., 1999), and closely ressambles spectra of several speech materials in other languages (Byrne et al., 1994; Wagener and Brand, 2005). The level of the masker noise was fixed at 70 dB SPL. Speech recognition thresholds for 50% correctly identified words (SRT_50_) were determined in 3 noise conditions: (i) quiet, (ii) ipsi-, (iii) contralateral noise (Garrett et al., 2020). For each 9 different combinations of noise condition, blocks of 20 sentences were presented per OLSA-BB, OLSA-LP and OLSA-HP condition. As a short training a test run of OLSA-BB and OLSA-HP with 20 sentences was performed before.

**Figure 2.**
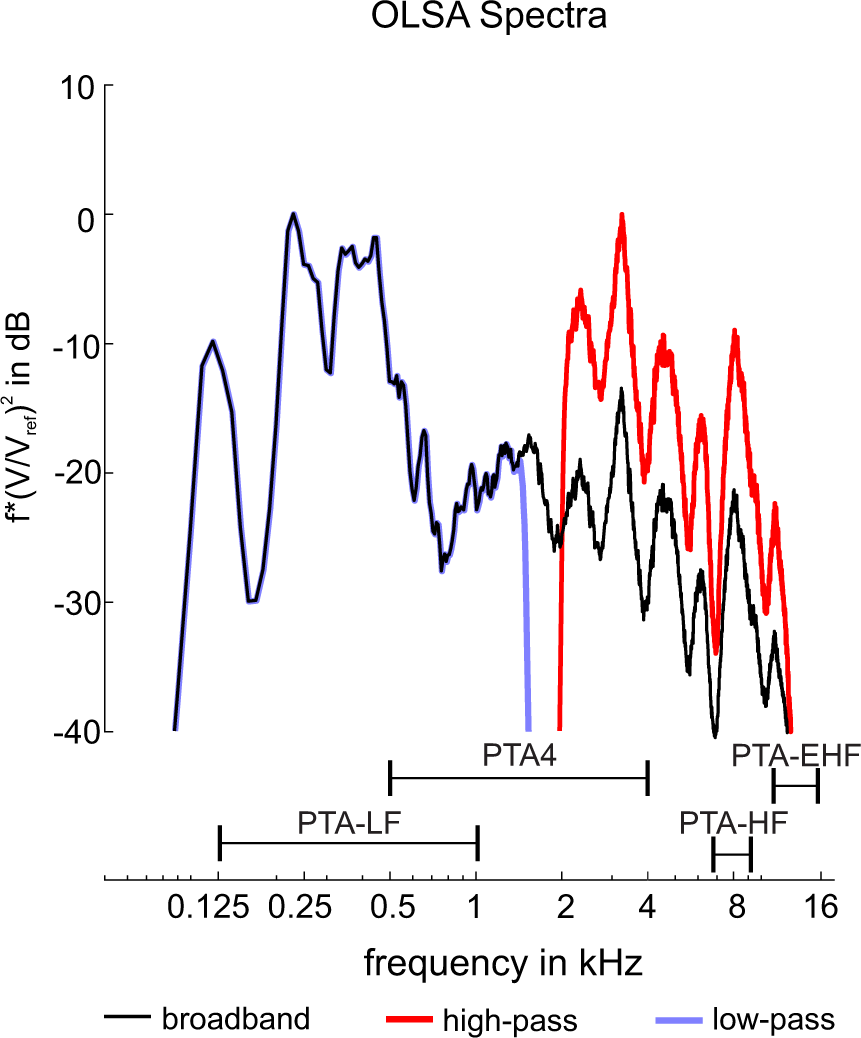
Power spectrum of the OLSA speech material (black curve), of high-pass filtered speech (HP-OLSA, red curve), and low-pass filtered speech (LP-OLSA, blue curve). Shown peak-normalized to 0 dB and 1/f-corrected. Three different filter version of OLSA stimuli demonstrated as power spectra.

### Pure-tone-normalized OLSA threshold

This study was particularly focused in factors beyond pure-tone thresholds and how they may relate to OLSA thresholds. In order to evaluate the role of these other factors, OLSA thresholds were quantitatively normalized for PTT of all available frequencies [0.125 - 16 kHz]. This correction was calculated independently for the quiet, ipsilateral-noise, and contralateral-noise condition by performing multivariate regression between all three different OLSA (BB, LP and HP) thresholds and the first five principle components (MatLab Version 2021b) of all audiometric thresholds, the latter in order to avoid over-fitting. The principal component analysis (PCA) was performed by employing a singular-value decomposition algorithm. Together, the first five PCA components capture 93% of the variations in audiometric thresholds. OLSA predictors for each individual subject were derived by evaluating the linear regression model using these first five components of the pure tone thresholds of each participant.

In order to evaluate the residual speech comprehension performance, we subtracted the three OLSA threshold predictions from the three measured OLSA thresholds and averaged them. This average value will be referred to as “PT-normalized OLSA threshold” or “PNOT”. Based on PNOT, the cohort was divided into three equally sized groups: with “good”, “normal” and “poor” speech performance. For details of group distribution, see results. We verified that this data-driven approach resulted in groups with matched average pure-tone audiometry thresholds within +-4.5 dB for the four PT frequency ranges (PTA-LF, PTA4, PTA-HF, PTA-EHF, see also Tab. 3).

**Table 3.** Frequencies of the first four formants (F1 - F4) for each individual vowel sound, measured in Hz. Vowel pair /o/ - /u/ had similar formants, only F1 was different. In vowel pair /i/ - /y/, /y/ had reduced formant frequencies F2, F3 and F4. Notably, F2 of /o/ - /u/ was located below the supposed phase- locking limit (∼1.5 kHz) while F2 of /i/ - /y/ was located above.

|  | /o/ | /u/ | /i/ | /y/ |
| --- | --- | --- | --- | --- |
| F1 | 380 | 200 | 350 | 350 |
| F2 | 750 | 750 | 1950 | 1780 |
| F3 | 3500 | 3500 | 2800 | 2100 |
| F4 | 10000 | 10000 | 4000 | 3000 |

### Stimuli for phoneme discrimination

The stimuli used for the phoneme discrimination task were computer-generated from actual recordings from a male speaker, using analysis-re-synthesis as implemented in the WORLD vocoder (Morise et al., 2016). Their average fundamental frequency **(F0)** was set to 116 Hz during synthesis in order to match the average F0 of the speaker of the OLSA sentences. A total of eight phonemes, two pairs of steady-state vowels and two pairs of consonant-vowel syllables were used as stimuli.

The vowels pairs /o/ (like in *oder*, “or”) and /u/ (like in *Du*, “you” in German) differed by their first formant (F1, see Tab.2), located well *below* the supposed phase-locking limit (PLL) in humans (∼1.5 kHz). They were synthesized with a 30-ms raised cosine ramp at onset and offset, and had a total duration of ∼414 ms (corresponding to 48 F0 cycles). Similarly, the /du/-/bu/ syllable pair only differed at frequencies below the PLL, and only within the first 100 ms. The following 371 ms were exactly identical between the two syllables. In addition, the vowel segment of this syllable pair, /u/, was identical to the isolated steady state /u/ used in the /o-u/ vowel pair, except that it was trimmed such that the overall duration of the syllables was 471 ms.

The vowel pairs /i/ (like in *die*) and /y/ (like in *üben*) only differed in their second and third formants (F2 and F3), which were *above* the PLL (see Tab. 3). As a result, it is expected that encoding of this /i/-/y/ contrast could not rely on temporal fine structure but on envelope coding.

These /i/-/y/ vowel contrast had the same duration and ramps as the other vowel pair presented above. The /di/-/bi/ syllable pair was also built to only differ in frequencies above the PLL, and within the spectral power of the first 100 ms. Again, the /i/ from these syllables was identical in spectral shape to the /i/ used in the vowel pair /i/-/y/.

All stimuli were then spectrally tilted to ensure similar signal-to-noise ratio above and below the PLL when presented in the speech-shape-noise used in the OLSA task.

From each of the four stimulus pairs, a 9-step continuum was generated by gradually modifying the formant(s) frequencies on a log-frequency scale. Following piloting, two contrasts were selected for each pair: a large contrast and a small contrast. Given the large inter-individual variability observed during piloting, this selection aimed to ensure that floor or ceiling effects would be avoided for at least one of these contrast magnitudes.

The stimuli were equalized such that the average level of the stimuli belonging to a given continuum was the same for all pairs, and was adjusted to 60 dB SPL (L_eq_). However, minor level fluctuations within a continuum were preserved to ensure that the level of the formants that remained identical throughout the continuum were not affected. Calibration was performed using a BK Type 4157 Microphone Brüel & Kjær (Böblinger Str. 13, 71229 Leonberg, Germany) in combination with an artificial ear with a volume of 1cm^3^ and a 20 s integration time.

### Behavioral phoneme discrimination task

The phoneme discrimination between the pairs (/o/-/u/, /i/-/y/, /du/-/bu/, /di/-/bi/) was measured using a three-alternative forced choice **(3AFC)** paradigm. For each phoneme pair, we measured two difficulty levels, an easy and a difficult difference. We quantified the difference in nine levels which were tested in pilot experiments from which two pairs, one easy and one difficult were selected for the psychoacoustic session. The differences in difficulty level e.g. for /du/-/bu/ in the easy condition was 8 out of 9, while in the difficult condition the difference was only 4 out of the 9. The respective other level differences for the difficult and easy condition were: for /di/-/bi/ 8 and 4, for /o/-/u/ 4 and 2 and for /i/-/y/ 3 and 1. Together with the three different noise conditions (quiet, ipsilateral, contralateral noise) we acquired data from a total of 6 conditions. Each condition was repeated nine times, producing a total of 54 trials. The noise was the same speech-shaped noise used during the OLSA measurement and presented at a 0 dB SNR.

To minimize learning effects, conditions were randomly reordered at the beginning of the measurement. As a short training a test run of 3 trials of phoneme discrimination of the four syllable pairs were performed before.

The right ear was used to test each syllable pair with the identical transducers as used for the OLSA test. Before each condition of the test, the participants were given four training trials with visual feedback. However, responses from this training were not included in data analysis. During the main test, participants did not receive any feedback on the correctness of their responses. Along with the responses, reaction time was recorded, even though not being in the main focus of the task.

### Statistical analysis

To test for the significance of group differences statistical tests for non-normally distributed data were applied. ABR wave amplitudes and latencies were compared by One-Way ANOVA for group differences. Resulting p-values smaller than the criterion of α = 0.05 were considered as statistically significant. The correlation of two measurement parameters was verified upon the Pearson Correlation Coefficient (r). ASSR response amplitudes (µV) were compared by Mann-Whitney-U tests between good and poor performers, between poor and standard performers, and between standard and good performers (1-sided hypothesis).

To test if syllable discrimination performance was different between the groups categorized for higher (good performers), lower (poor performers), or standard speech recognition performance based on PNOT, the score of percent-correct answers was compared by Mann-Whitney-U tests between good and poor performers, between poor and standard performers, and between standard and good performers (1-sided hypothesis: The group of poor/standard performers contains more participants with low-percentage correct scores relative to the standard/good performers). A resulting p-value equal or smaller than α = 0.05 was considered statistically significant and noted as asterisk in the respective figure panel (Fig. 3). A p-value smaller 0.1 was noted as asterisk in brackets to inform for a trend in the distribution, though not reaching statistical significance. Statistical comparisons were done for the percent correct scores obtained for the “difficult” discrimination task (small spectral and temporal syllable contrast) and for the scores obtained for the “easy” discrimination task (larger spectral and temporal syllable contrast).

**Figure 3.**
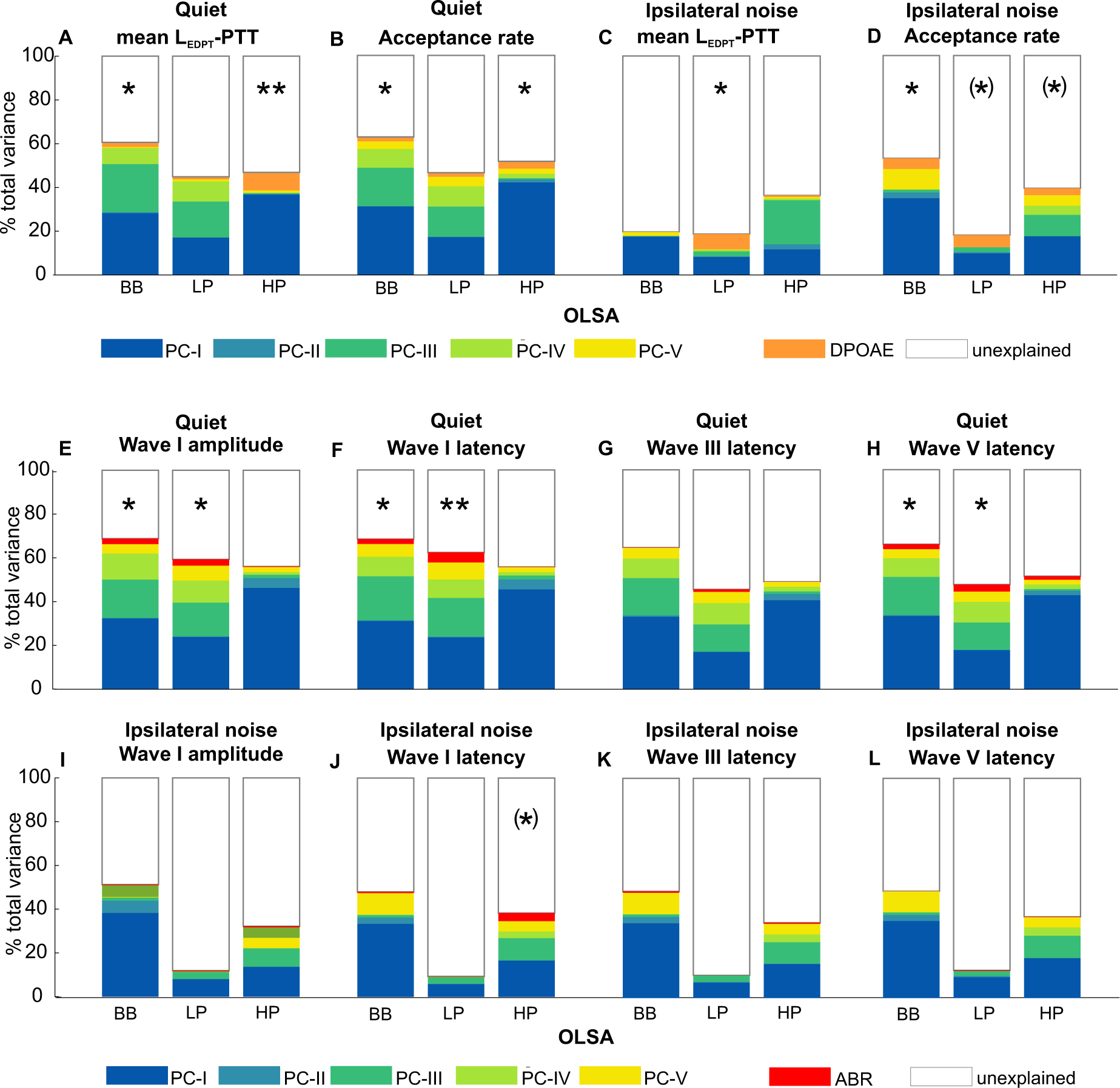
Posthoc analysis of OLSA variance based on a linear mixed model (A-D) DPOAE: Ledpt-PTT contributing 2% to OLSA bb and 8.3% to OLSA hp in quiet (A) and 7% (lp) in ipsilateral noise (C). DPOAE acceptance rates (B, D) show that the number of valid LEDPT measurements contributing (B) 2% OLSA bb and 3.3% OLSA hp in quiet and (D) 4,8% OLSA bb in ipsilateral noise. (E-L) ABR wave I amplitude (E, I) and latencies for wave I (F, J), II (G, K) and V (H, L) contributing to SRT variance (percentages see text). Significance levels based on permutation testing: * p<0.05 and ** p<0.01. Tendencies with p<0.1 are marked with a dot.

### Variance analysis

Analysis of variance for speech perception thresholds beyond pure tone thresholds was performed by least square multivariate linear fitting of the 5 added PCs derived from pure tone thresholds and one additional observable which was tested for its contribution to total speech comprehension variance. To ensure uniqueness of the multivariate linear model, we first removed all linear correlation between the 5 PCs and the tested observable. This can be understood as removing the influence of PTT on ABR wave amplitude or latency or other parameters like L_EDPT_ thresholds, ASSR amplitudes or phoneme discrimination. An inherent risk of the increase of dimensions of the regression model is the possibility for over-fitting. In order to eliminate this effect, we compared the observed increase in explainable variance in the observed 6-dimensional model to the variance of 10,000 pseudo-models in which we randomly shuffled the additional observable before fitting the model. This gives us a reliable estimate of what gain in explained variance was achieved based on chance. The results are presented as stacked bar diagrams showing the percentage of variance which could be attributed to each of the observables (Tab. 4). To better illustrate the magnitude of the effect, we additionally computed the standard deviation in the unit of the speech reception thresholds **(SRT)** [dB] which can be attributed to the new observable, by taking the ratio of the variance of the observable to the overall variance, and multiplying with the SD of the sample in [dB] (Fig. 3). However, during this computation we assumed normal distribution of the 5 PCs and the additional tested observable, an assumption which is not required for the statistical evaluation based on permutation analysis.

**Table 4.** Posthoc analysis of OLSA variance based on a linear mixed model. The left most column contains the parameters (LEDPT, Number of DPOAE, ABR amplitude wave I and latencies waves I, II, III, V, VI, ASSR, psychoacoustic results on phoneme discrimination) which were tested for contributions to explain SRT beyond PTTs. The second column represents the respective size of sub-samples qualified for this analysis. The 5 result columns are all split for SRT in the “bb” = broadband, “lp” = low pass and “hp” = high pass filtered conditions. The result columns “sign of correlation”, “explained total variance in %”, “explained standard deviation (std) in dB”, “p-value” and “total std dB” were are all derived from 10.000 boot straps and provide significance based on the probability that the parameter under test does not contribute. The green shading of the cells in the “p-value” column reflect significance levels: in light green = trends (p<0.1), mid tone green = 5% significance level and dark green = highly significant p<0.01.

|  |  |  | sign of correlation |  |  | Explained total Variance in % |  |  | Explain std dB |  |  | p value |  |  | Total std dB |  |  |
| --- | --- | --- | --- | --- | --- | --- | --- | --- | --- | --- | --- | --- | --- | --- | --- | --- | --- |
|  |  | N | bb | lp | hp | bb | lp | hp | bb | lp | hp | bb | lp | hp | bb | lp | hp |
| LEDPT-PTT | Quiet | 81 | 1 | 1 | 1 | 2.0 | 1.2 | 8.3 | 0.8 | 0.7 | 4.8 | 0.033 | 0.161 | 0.001 | 5.5 | 6.6 | 16.6 |
|  | Ipsi | 56 | 1 | 1 | 1 | 0.1 | 7.0 | 0.7 | 0.0 | 0.3 | 0.5 | 0.783 | 0.037 | 0.519 | 1.1 | 1.3 | 6.0 |
| Number of DPOAE | Quiet | 88 | -1 | -1 | -1 | 2.0 | 1.8 | 3.3 | 0.8 | 0.9 | 3.2 | 0.037 | 0.105 | 0.019 | 5.9 | 6.7 | 17.8 |
|  | Ipsi | 63 | 1 | 1 | 1 | 4.8 | 5.5 | 3.1 | 0.3 | 0.3 | 1.1 | 0.023 | 0.051 | 0.079 | 1.4 | 1.3 | 6.1 |
| ABR amp I | Quiet | 76 | -1 | -1 | 1 | 2.5 | 2.8 | 0.2 | 0.9 | 1.0 | 0.8 | 0.022 | 0.032 | 0.561 | 5.4 | 6.2 | 16.7 |
|  | Ipsi | 56 | -1 | -1 | 1 | 0.3 | 0.3 | 0.5 | 0.1 | 0.1 | 0.4 | 0.634 | 0.671 | 0.525 | 1.5 | 1.3 | 6.0 |
| ABR lat I | Quiet | 80 | 1 | 1 | -1 | 2.2 | 4.6 | 0.1 | 0.8 | 1.3 | 0.5 | 0.027 | 0.004 | 0.727 | 5.6 | 6.3 | 16.5 |
|  | Ipsi | 60 | -1 | 1 | -1 | 0.5 | 0.1 | 3.7 | 0.1 | 0.0 | 1.2 | 0.491 | 0.790 | 0.063 | 1.5 | 1.2 | 6.0 |
| ABR lat II | Quiet | 62 | 1 | 1 | 1 | 4.0 | 4.3 | 2.1 | 1.0 | 1.2 | 2.3 | 0.018 | 0.053 | 0.128 | 5.2 | 5.8 | 16.1 |
|  | Ipsi | 47 | -1 | 1 | -1 | 0.3 | 0.2 | 3.6 | 0.1 | 0.1 | 1.1 | 0.675 | 0.745 | 0.110 | 1.6 | 1.2 | 5.9 |
| ABR lat III | Quiet | 84 | 1 | 1 | -1 | 0.1 | 1.1 | 0.0 | 0.2 | 0.7 | 0.3 | 0.612 | 0.211 | 0.811 | 5.7 | 6.7 | 17.1 |
|  | Ipsi | 61 | 1 | -1 | -1 | 0.7 | 0.0 | 0.5 | 0.1 | 0.0 | 0.4 | 0.427 | 0.908 | 0.487 | 1.4 | 1.2 | 6.0 |
| ABR lat V | Quiet | 86 | 1 | 1 | 1 | 2.2 | 3.2 | 1.6 | 0.8 | 1.2 | 2.2 | 0.024 | 0.030 | 0.112 | 5.6 | 6.6 | 17.6 |
|  | Ipsi | 62 | 1 | -1 | -1 | 0.0 | 0.3 | 0.1 | 0.0 | 0.1 | 0.2 | 0.858 | 0.667 | 0.720 | 1.4 | 1.3 | 6.2 |
| ABR lat VI | Quiet | 78 | 1 | 1 | 1 | 1.0 | 2.0 | 4.2 | 0.5 | 0.9 | 3.3 | 0.153 | 0.105 | 0.011 | 5.4 | 6.7 | 16.2 |
|  | Ipsi | 58 | -1 | -1 | -1 | 0.1 | 0.4 | 0.3 | 0.1 | 0.1 | 0.3 | 0.726 | 0.613 | 0.650 | 1.4 | 1.3 | 5.9 |
| ASSR | Quiet | 74 | 1 | 1 | 1 | 0.4 | 0.1 | 0.9 | 0.4 | 0.2 | 1.6 | 0.387 | 0.749 | 0.273 | 5.8 | 6.8 | 16.3 |
|  | Ipsi | 55 | 1 | 1 | 1 | 7.0 | 4.3 | 2.0 | 0.4 | 0.3 | 0.9 | 0.012 | 0.109 | 0.219 | 1.4 | 1.3 | 6.2 |
| ASSR 4k | Quiet | 65 | 1 | -1 | 1 | 0.0 | 0.7 | 0.5 | 0.1 | 0.6 | 1.1 | 0.875 | 0.336 | 0.447 | 5.9 | 6.9 | 15.1 |
|  | Ipsi | 49 | 1 | 1 | -1 | 0.3 | 0.9 | 0.1 | 0.1 | 0.1 | 0.2 | 0.612 | 0.487 | 0.805 | 1.2 | 1.2 | 5.9 |
| ASSR 6k | Quiet | 68 | 1 | 1 | 1 | 1.2 | 0.5 | 0.2 | 0.6 | 0.5 | 0.7 | 0.150 | 0.407 | 0.632 | 5.2 | 6.7 | 17.7 |
|  | Ipsi | 50 | 1 | 1 | 1 | 3.8 | 0.4 | 3.3 | 0.3 | 0.1 | 1.1 | 0.085 | 0.661 | 0.147 | 1.4 | 1.3 | 6.3 |
| di bi easy | Quiet | 73 | -1 | -1 | 1 | 2.3 | 1.7 | 0.2 | 0.9 | 0.9 | 0.8 | 0.038 | 0.146 | 0.603 | 6.1 | 7.0 | 18.2 |
|  | Ipsi | 51 | -1 | 1 | -1 | 0.1 | 0.1 | 0.2 | 0.0 | 0.0 | 0.3 | 0.761 | 0.842 | 0.667 | 1.4 | 1.3 | 6.2 |
| ou difficult | Quiet | 88 | -1 | -1 | -1 | 2.0 | 0.0 | 0.2 | 0.9 | 0.1 | 0.7 | 0.035 | 0.863 | 0.603 | 6.0 | 6.7 | 17.8 |
|  | Ipsi | 62 | -1 | -1 | -1 | 0.4 | 1.7 | 1.6 | 0.1 | 0.2 | 0.8 | 0.533 | 0.289 | 0.216 | 1.4 | 1.3 | 6.2 |
| ou easy | Quiet | 88 | -1 | -1 | -1 | 2.0 | 0.7 | 1.2 | 0.8 | 0.5 | 1.9 | 0.039 | 0.319 | 0.169 | 6.0 | 6.7 | 17.8 |
|  | Ipsi | 62 | 1 | 1 | -1 | 0.1 | 0.0 | 0.7 | 0.0 | 0.0 | 0.5 | 0.765 | 0.899 | 0.406 | 1.4 | 1.3 | 6.2 |
| iy difficult | Quiet | 88 | -1 | -1 | -1 | 0.6 | 0.5 | 0.5 | 0.5 | 0.5 | 1.2 | 0.256 | 0.384 | 0.398 | 6.0 | 6.7 | 17.8 |
|  | Ipsi | 62 | -1 | 1 | -1 | 4.1 | 0.1 | 2.6 | 0.3 | 0.0 | 1.0 | 0.041 | 0.748 | 0.106 | 1.4 | 1.3 | 6.2 |
| iy easy | Quiet | 88 | 1 | 1 | -1 | 0.0 | 0.1 | 0.5 | 0.1 | 0.2 | 1.3 | 0.892 | 0.664 | 0.359 | 6.0 | 6.7 | 17.8 |
|  | Ipsi | 62 | -1 | 1 | -1 | 1.3 | 0.3 | 1.0 | 0.2 | 0.1 | 0.6 | 0.248 | 0.680 | 0.334 | 1.4 | 1.3 | 6.2 |
| UCL 250 | Quiet | 55 | -1 | 1 | 1 | 0.0 | 0.0 | 0.1 | 0.1 | 0.1 | 0.6 | 0.853 | 0.839 | 0.729 | 6.2 | 7.2 | 15.9 |
| UCL 250 | Ipsi | 42 | 1 | 1 | -1 | 0.0 | 10.1 | 0.7 | 0.0 | 0.4 | 0.5 | 0.969 | 0.024 | 0.497 | 1.5 | 1.2 | 6.2 |
| UCL 500 | Quiet | 69 | -1 | 1 | 1 | 0.0 | 0.0 | 1.5 | 0.1 | 0.1 | 2.1 | 0.863 | 0.839 | 0.182 | 6.2 | 7.2 | 17.4 |
| UCL 500 | Ipsi | 52 | 1 | 1 | -1 | 0.3 | 7.6 | 0.0 | 0.1 | 0.4 | 0.1 | 0.619 | 0.036 | 0.906 | 1.5 | 1.3 | 6.5 |
| UCL 1000 | Quiet | 72 | -1 | -1 | 1 | 0.2 | 0.1 | 1.9 | 0.3 | 0.2 | 2.4 | 0.544 | 0.726 | 0.117 | 6.2 | 7.1 | 17.8 |
| UCL 1000 | Ipsi | 53 | 1 | 1 | -1 | 0.4 | 8.6 | 0.0 | 0.1 | 0.4 | 0.1 | 0.533 | 0.022 | 0.895 | 1.5 | 1.3 | 6.4 |
| UCL2000 | Quiet | 80 | -1 | -1 | 1 | 0.0 | 0.1 | 2.1 | 0.0 | 0.2 | 2.6 | 0.915 | 0.776 | 0.078 | 6.0 | 6.8 | 17.8 |
| UCL2000 | Ipsi | 56 | 1 | 1 | -1 | 0.1 | 4.4 | 0.2 | 0.1 | 0.3 | 0.2 | 0.718 | 0.099 | 0.696 | 1.4 | 1.3 | 6.3 |
| UCL4000 | Quiet | 77 | 1 | 1 | 1 | 0.1 | 0.2 | 1.2 | 0.2 | 0.3 | 1.9 | 0.692 | 0.644 | 0.209 | 5.9 | 6.9 | 17.4 |
| UCL4000 | Ipsi | 56 | 1 | 1 | -1 | 0.3 | 4.2 | 0.1 | 0.1 | 0.3 | 0.2 | 0.572 | 0.098 | 0.796 | 1.4 | 1.3 | 6.3 |
| UCL 6000 | Quiet | 42 | 1 | -1 | 1 | 0.1 | 0.5 | 1.0 | 0.2 | 0.6 | 1.9 | 0.782 | 0.616 | 0.417 | 7.1 | 8.4 | 19.0 |
| UCL 6000 | Ipsi | 25 | -1 | -1 | -1 | 1.2 | 1.4 | 0.0 | 0.2 | 0.2 | 0.0 | 0.455 | 0.515 | 0.947 | 1.4 | 1.4 | 5.9 |

### Data Distributions

If not indicated otherwise, data are presented as group mean and standard deviation (SD) for the number of participants or ears (n) specified in the figure legends. Syllable discrimination performance (% correct) was classified in histograms with logarithmic class sizes for visualization of the different performances of poor, good, and standard speech recognition.

## RESULTS

We included 89 participants evenly distributed across three age-groups of young (18-29 years, n = 29), middle-aged (30-55 years, n = 32) and older (56-76 years, n = 28) participants as described in methods. A five-grade custom questionnaire for subjective self-evaluation of hearing performance (excellent, very good, good, moderate, bad) in different conversation-situations was conducted, and three psychometric tests were performed to exclude any confounding severe psychiatric factors, such as depression or dementia onset. The analysis of the results of BDI, GDS, and MMSE across age (methods, Fig. 1) revealed two young participants, one middle-aged participant, and one older participant who each scored higher than normal in the BDI (> 20) and GDS (>6) (Fig. 1A,B, red line). These individuals are specifically highlighted throughout the following approaches.

### Correlation of pure-tone thresholds with age

PTT were collected for frequencies between 0.125 to 16 kHz as described in methods. Comparison of the three age groups revealed group differences that shifted towards significantly elevated thresholds above 8 kHz, particularly prominent for EHF thresholds obtained between 11.2 and 16 kHz (Fig. 4A), as was also observed in previous studies (Osterhammel and Osterhammel, 1979; Prendergast et al., 2020; Carcagno and Plack, 2022; Marcher-Rorsted et al., 2022).

**Figure 4.**
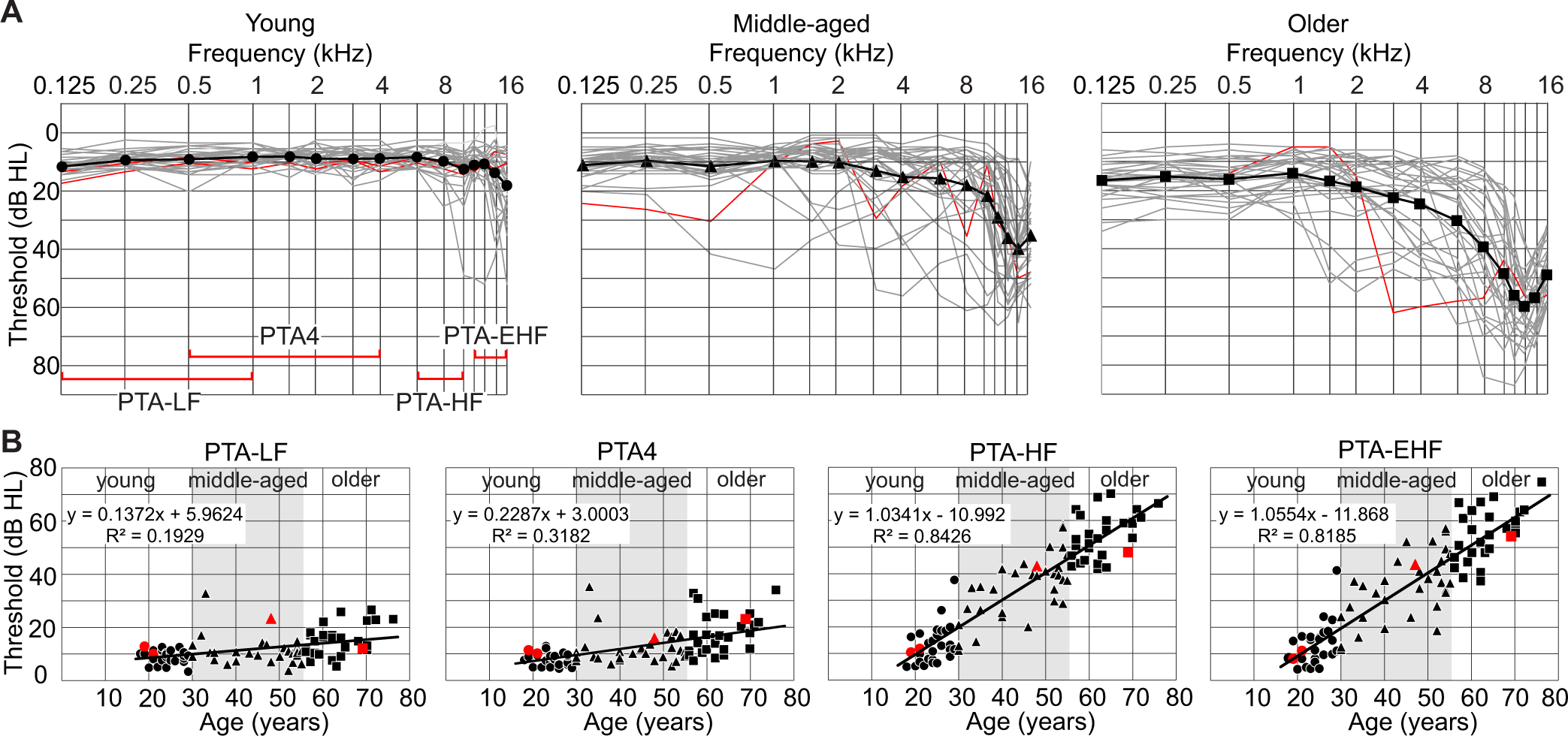
Elevated hearing thresholds correlate with age, in particular at high frequencies. (A) Pure tone thresholds for the three age groups young (left), middle-aged (center) and elderly (right) measured for tone frequencies between 0.125 kHz and 16 kHz, assigned to four different frequency ranges low frequencies “PTA-LF” [0.125 - 1 kHz], “PTA4” [0.5 - 4 kHz], high frequencies “PTA-HF” [6 – 10 kHz] and extended high frequencies “PTA-EHF” [11.2-16 kHz], illustrated on the abscissa of the leftmost audiogram. The group mean thresholds are plotted in black (young group with circles, middle aged with triangles and elderly with squares).(B) Scatterplots for individual hearing thresholds as a function of age, split for the four frequency ranges. The shaded area delineates the age range of the middle-aged group. Circles: young group; triangles: middle-aged group; squares, older group; red symbols: subjects outside the normal range in both the BDI and GDS tests. P-Value (Pearson’s correlation) p(PTA-LF)= 0.000016; p(PTA4) <0.00001; p(PTA-HF) <0.00001, p(PTA-EHF) <0.00001.

All four PTAs, averaged over different frequency ranges, correlated significantly with age (Fig. 4B, PTA-LF: p= 0.000016, R^2^=0.1929; PTA4: p <0.00001, R^2^=0.3182; PTA-HF: p<0.00001, R^2^=0.8426; PTA-EHF: p<0.00001, R^2^ =0.8185). The slope of the regression lines (see Fig. 4B, R^2^ values) is much steeper if correlation is computed with respect to the HF and EHF averages, with considerably higher r^2^ values than for the lower frequency averages (0.84 and 0.81 as compared to 0.19 and 0.32 for PTA-LF and PTA4, Fig. 4B), supporting the notion that age-dependent hearing loss is predominantly an increasing loss of high frequency hearing. The four subjects who were outside the normal levels in the BDI and GDS tests (Fig. 1) showed no abnormalities in the PTT over the entire frequency range (Fig. 4A, B, red symbols).

### SRT in quiet, ipsi- and contralateral noise depends on PTA-threshold and age

SRT were determined for OLSA-BB, OLSA-LP, and OLSA-HP (Fig. 5A-C) and plotted as a function of age. We observed that SRT in quiet were significantly positively correlated with age, with the strongest dependence for the high-pass filtered condition (Fig. 5C, A: R^2^= 0.2562, p= < .00001; B: R^2^= 0.1376, p= 0.000345; C: R^2^= 0.3631, p= < .00001; D: R^2^= 0.186, p= 0.000417; E: R^2^= 0.0669, p= 0.040715; F: R^2^= 0.1292, p= 0.003807; G: R^2^= 0.2632, p= 0.000017; H: R^2^= 0.1732, p= 0.000692; I: R^2^= 0.4306, p= < .00001). SRT under speech-shaped ipsilateral noise were also significantly correlated with age, but the dependence on age was markedly reduced (Fig. 5D). When OLSA stimuli were low- or high-pass filtered (Fig. 5E, F), SRT in ipsilateral noise was elevated, and the respective slopes were shallower than the slopes for the speech-in-quiet condition. In contrast to ipsilateral noise, contralateral speech-shaped noise yielded similar OLSA thresholds and its age-dependence as those obtained in quiet in any of the OLSA filter conditions (Fig. 5G, H, I).

**Figure 5.**
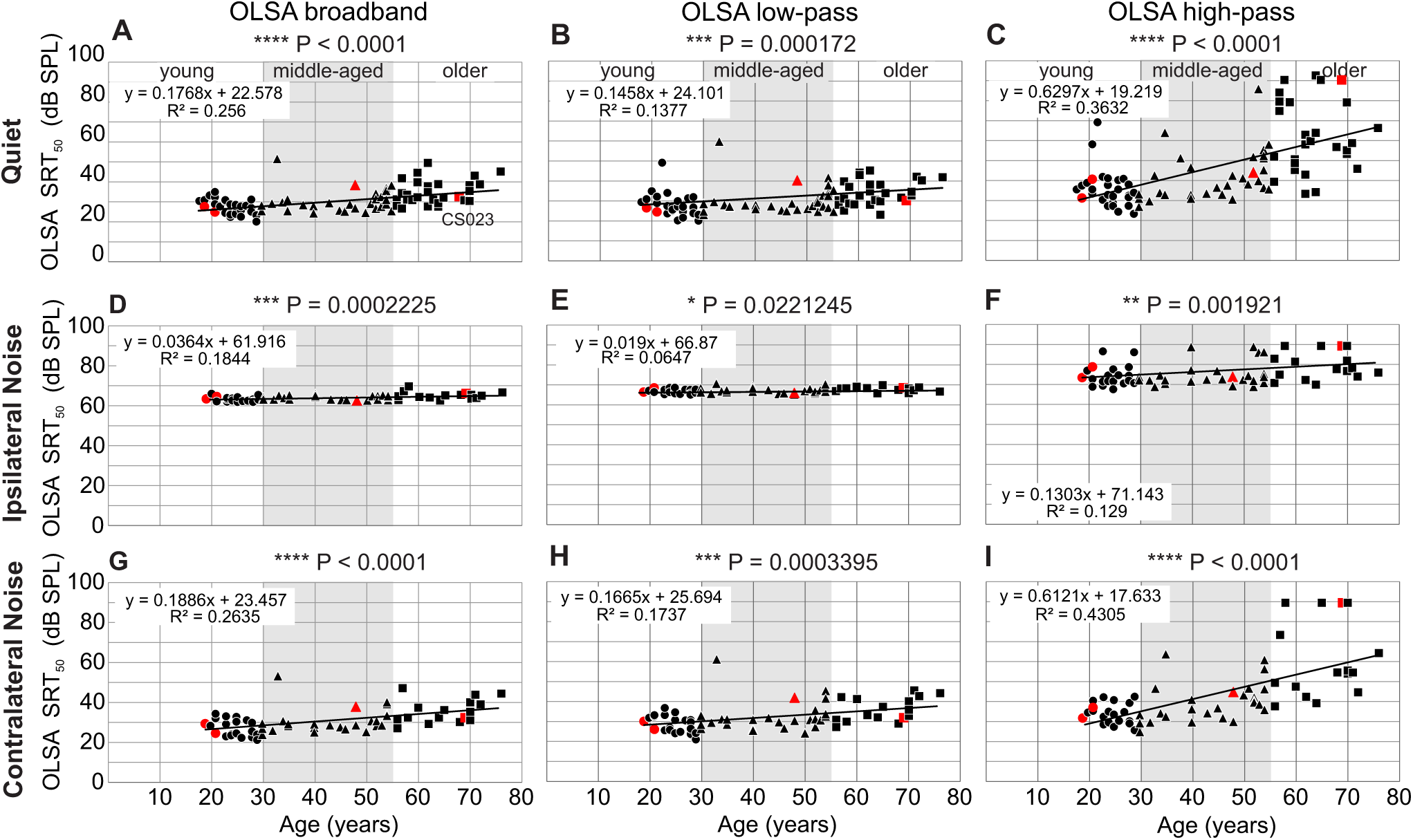
Influence of noise on OLSA speech recognition threshold SRT50, determined with different bandwidths. Noise conditions in rows: A-C quiet, D-F ipsilateral noise, G-I contralateral noise. Bandwidth of OLSA in columns: A, D, G provide results for broadband OLSA stimuli, B, E, H provide results for low-pass [< 1500 Hz] filtered OLSA stimuli, C, F, I provide results for high-pass [>1500 Hz] filtered OLSA stimuli. Each provides the respective OLSA SRTs as a function of age. Circles: young group; triangles: middle-aged group; squares, older group; red symbols: subjects outside the normal range in both the BDI and GDS tests. P, see inset; y, r2 for trendline and significance of corresponding P-value for Pearson Correlation Coefficient.

In conclusion, SRTs significantly depend on age in all conditions. The four subjects who exceeded the critical scores in the BDI and GDS tests (Fig. 1) performed normal for all OLSA conditions (Fig. 5, red dots).

Fig. 6 shows the dependence of all OLSA SRT on their corresponding pure-tone averages — i.e., dependence of OLSA-BB on PTA4 (Fig. 6A, D, G), dependence of OLSA-LP on PTA-LF (Fig. 6B, E, H), and dependence of OLSA-HP on PTA-EHF (Fig. 6C, F, I). In all nine comparisons, OLSA SRT significantly depended on the corresponding PTA measure (Fig. 6, A: R^2^= 0.5363, p= < .00001; B: R^2^= 0.4574, p= < .00001; C: R^2^= 0.378, p= < .00001; D: R^2^= 0.3111, p= < .00001; E: R^2^= 0.064, p= 0.045525; F: R^2^= 0.1748, p= 0.00065; G: R^2^= 0.5799, p= 0.00065; H: R^2^= 0.6254, p= < .00001; I: R^2^= 0.5032, p= < .00001), with stronger scatter for the dependence of OLSA-HP SRT on PTA-EHF in all three noise conditions tested (Fig. 6C, F, I). In quiet as well as in contralateral noise, OLSA-BB depended on PTA4 with a slope of the regression line of 0.63 and 0.65, and R^2^=0.54 and 0.58 (Fig. 6A, G). Interestingly, similar correlation coefficients were also obtained for the dependence of OLSA-LP on PTA-LF (R^2^=0.46 and 0.63; Fig. 6B, H) and for the dependence of OLSA-HP on PTA-EHF (R^2^=0.38 and 0.50; Fig. 6C, I). The comparatively higher scatter in the high-frequency conditions along with lower R^2^ figures (Fig. 6C) is a result of the mismatch between the OLSA filter condition passing frequencies above 1.5 kHz (chosen to match the PLL) and the frequency content of the PTA-EHF.

**Figure 6.**
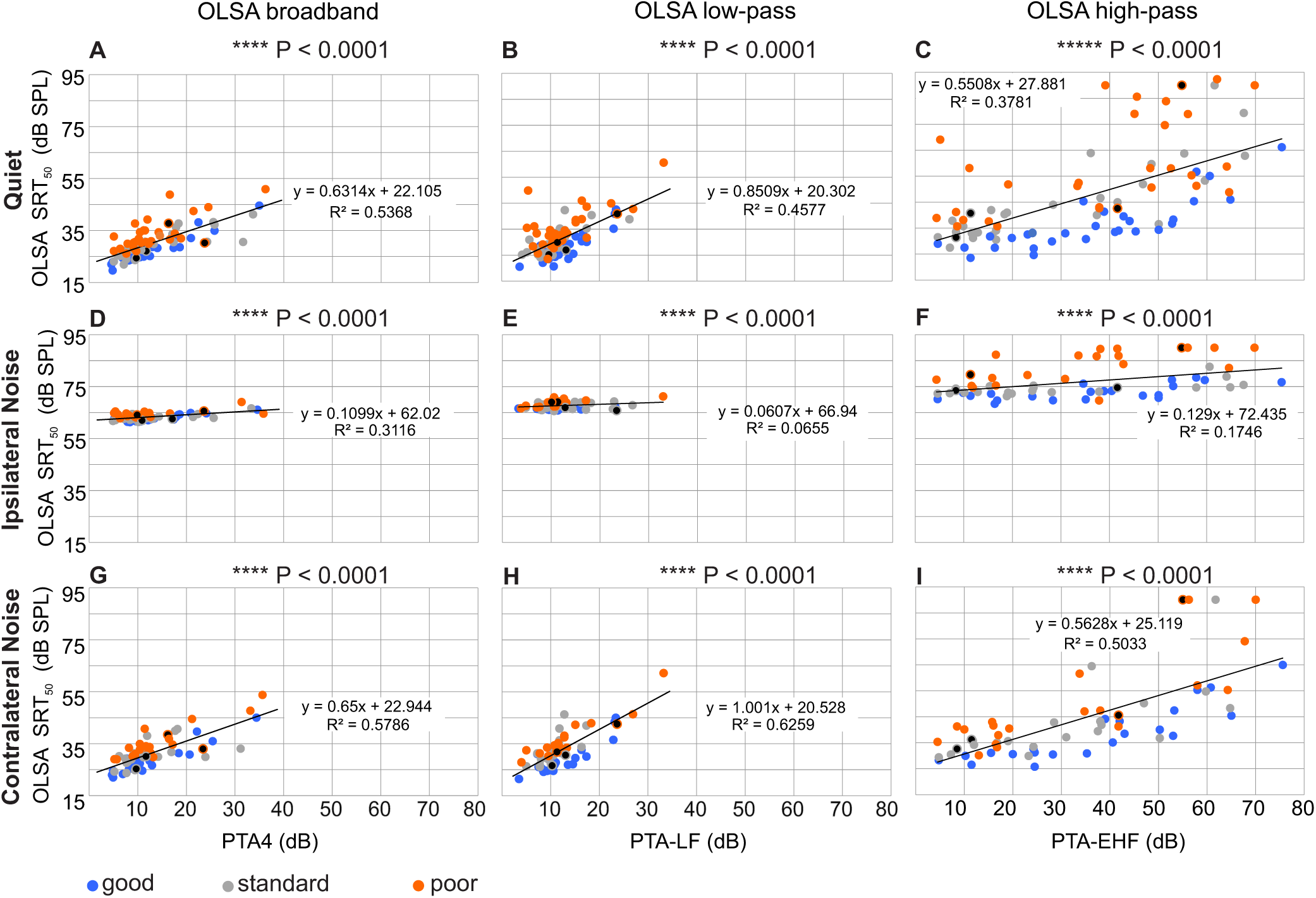
OLSA speech recognition threshold SRT50 (dB SPL, y-axes) plotted for three bandwidths of OLSA stimuli. Broadband (left column), low-pass (middle column) and high-pass (right column) are shown as functions (x-axes) of PTA4 (left column), PTA-LF (middle column), and PTA-EHF (right column). The top row of plots provides results obtained in quiet (n = 89), middle row under ipsilateral (n = 63) and bottom row under contralateral speech-shaped noise conditions (n = 63). Regression lines are plotted in black. including y-intersections and R² values. The different colors reflect the three speech comprehension groups: good (blue), standard (grey) and poor (orange). Note: none of the subjects who were outside the normal levels in the BDI and GDS tests (circled in black) fell into to good (blue) group.

For OLSA in ipsilateral noise, the correlation shows a clearly reduced slope of the regression lines in all frequency conditions. For OLSA-BB (Fig. 6D), the slope of approximately 0.1 dB/dB almost perfectly reproduced previous findings (Wardenga et al., 2015). For the low-pass condition (Fig. 6E), the slope drops to 0.06, and only slightly increases for the high-pass condition to 0.13 (Fig. 6F). Similar to the investigation of (Wardenga et al., 2015), we see a fairly homeostatic distribution around the regression line for hearing losses up to 40 dB, indicating that the part of speech-in-noise intelligibility which cannot be explained by pure-tone threshold in noise, is present starting from the age of 20 over the complete range and thus appears not to be a pathologic state which accumulates with age.

Interestingly, in all nine conditions shown in Fig. 5 and 6, the OLSA SRT correlated better (higher R^2^) with the patient’s PTA than with patient’s age, an effect which is very clear for speech in quiet and contralateral noise (Fig. 6A-C; G-I; mean R^2^ = 0.51 vs. 0.27), and less pronounced for speech in ipsilateral noise (Fig. 6D; OLSA-BB: R^2^ = 0.31 vs. 0.18), where it almost vanishes only for OLSA-LP vs. PTA-LF (Fig. 6E).

For *post hoc* analysis, we used a classification that was based on PTT-compensated OLSA (PNOT), as described in methods. The multivariate regression on OLSA thresholds returned R² = 0.49. The remaining variance of the speech intelligibility not explained by PTT suggests unknown additional pathogenic mechanisms or measurement uncertainty.

This approach was used to separate the cohort into three groups with a maximum spread in their speech comprehension (Fig. 6, see blue = “good” and orange = “poor” dots), as well as the “standard” group with comprehension performance between good and poor groups (Figure 6, grey dots).

As expected, participants with poor, standard, and good speech comprehension based on PNOT showed well-matched mean PTT (Tab. 5). Interestingly, poor and good performers showed almost the same mean age, whereas the standard PNOT-group was on average slightly younger and exhibited slightly but not significantly better PTA-EHF-thresholds: PNOT-quiet: p(age)=0.100; p(PTA-EHF)=0.150; PNOT-contra: p(age)=0.606; p(PTA-EHF)=0.529; PNOT-ipsi: p(age)=0.713; p(PTA-EHF)=0.547 (Tab.5). 77.8% of the participants with good, standard, and poor speech comprehension in contralateral noise were grouped into the same categories in the speech-in-quiet tasks. The same can be said for only 39.9% of participants from the speech comprehension in ipsilateral noise tasks. For future analyses we therefore consider the PNOT in quiet and PNOT in contralateral noise as one group that may have different characteristics from the PNOT in ipsilateral noise group.

**Table 5.** Pure tone threshold normalized speech comprehension thresholds SRT (PNOT) differentiated for three noise conditions. Sub-sample sizes, mean age, and standard error of the mean (SEM) for PTA4 and PTA-EHF thresholds (dB HL) are given for participants with good, standard and poor speech comprehension.

| Noise conditions | PNOT | N | mean age (SEM) | mean PTA4 (SEM) | mean PTA-EHF (SEM) |
| --- | --- | --- | --- | --- | --- |
| In quiet | good | 30 | 45.40 +- 2.87 | 12.33 +- 1.21 | 36.22 +- 3.24 |
|  | standard | 29 | 38.48 +- 3.10 | 13.41 +- 1.44 | 28.67 +- 3.73 |
|  | poor | 30 | 47.60 +- 3.25 | 13.40 +- 1.18 | 38.28 +- 3.73 |
| In contralateral noise | good | 21 | 45.52 +-3.67 | 12.69 +- 1.65 | 36.85 +- 4.30 |
|  | standard | 21 | 40.24 +- 3.44 | 12.36 +- 1.38 | 29.85 +- 4.09 |
|  | poor | 21 | 42.62 +- 4.04 | 14.57 +- 1.75 | 33.01 +- 4.70 |
| In ipsilateral noise | good | 21 | 45.24 +- 3.64 | 14.02 +- 1.49 | 36.00 +- 4.19 |
|  | standard | 21 | 41.05 +- 3.87 | 13.21 +- 1.53 | 29.42 +- 4.67 |
|  | poor | 21 | 42.10 +- 3.70 | 12.38 +- 1.78 | 34.30 +- 4.25 |

Interestingly, none of the four subjects who were outside the normal levels in the BDI and GDS tests (Fig. 1) were distributed in either the good PNOT- in quiet group or the good PNOT- in ipsi- noise group (Fig. 6, red dots).

Speech-in-quiet comprehension, used here essentially as a control, depended strongly on PTT with a slope of more than 0.6 dB/dB. When removing the effect of PTT, in OLSA-BB, 38.7% of the variance remained unexplained, corresponding to a SD of 3.7 dB in SRT. In contrast, broadband speech-in-noise comprehension depended on PTT with a slope of approximately 0.1 dB/dB, leaving 51.4% of variance unexplained when removing the effect of PTT, corresponding to a SD of 1.0 dB in SRT.

To explain remaining variance, we tested participants with good and poor speech performance based on the PNOT measures in either quiet, ipsilateral noise, or contralateral noise for relations to **(i)** subjective speech understanding, **(ii)** ASSR, **(iii)** precise measurements of the cochlear amplifier using short-pulsed DPOAEs, **(iv)** central suprathreshold auditory brainstem responses, **(v)** phoneme discrimination ability.

### Delta to pure tone normalized OLSA thresholds (PNOT) are a better indicator of self-assessed hearing ability than age

Subjective self-evaluation of language comprehension in different conversation situations was acquired with our custom questionnaire (see methods). We compared the self-evaluation of young, middle-aged, and older participants (Fig. 7A) as well in participants previously grouped by good, standard, and poor speech comprehension according to PNOT (Fig. 7B). We did find that the PNOT classification of participants was a better predictor of their subjective assessment of their hearing than their age. In particular, a higher percentage of the middle-aged and older participants rated themselves as hearing very well – comparable to the young population – when sub-division was assessed according to age (Fig. 7A, n(young)=29, n(middle-aged)=32, n(older)=28, p = 0.50, one-sided Fisher Exact Probability Test for “very good” and “good” assessments). However, when patients were instead classified as having good, standard, or poor speech comprehension by the PNOT in quiet, we found that the expected self-rated decrease in hearing ability with age was much more congruent with real hearing performance, though not reaching statistical significance (Fig. 7B, n(good)=30, n(standard)=29, n(poor)=30, p = 0,10, one-sided Fisher Probability Test for “very good” and “good” assessments), with no differences when grouped for PNOT in ipsilateral noise (n(good, standard, poor)=21, p = 0.50) but again more congruent for contralateral noise (n(good, standard, poor)=21, p = 0.02, one-sided Fisher Exact Probability Test for “very good” and “good” assessment, not shown in Fig. 7).

**Figure 7.**
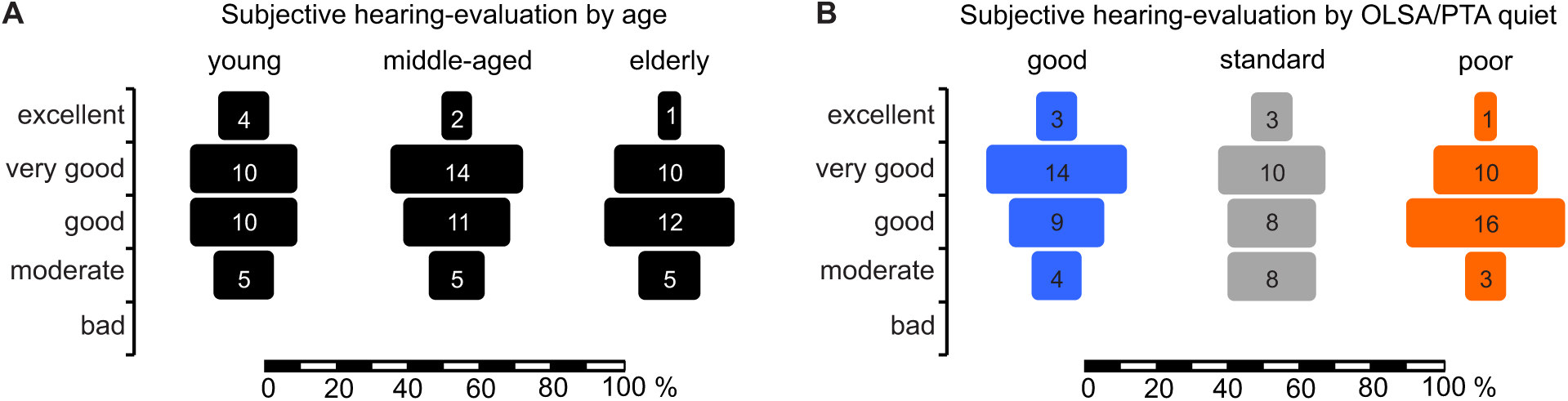
Subjective hearing evaluation by age and speech comprehension. (A) grouping by age and (B) grouping by objective speech comprehension performance based on OLSA thresholds corrected for by PNOT. Y-axis: Self-rated hearing ability in percent. X-axis: percentage of all survey responses given by all participants in each age group (A) and in each PNOT-group (B). Participants were asked to rate their hearing as excellent, very good, good, moderate or bad (see y-axis labels).

### Supra-threshold ASSR responses are not reflected in PNOT grouping

To address a potential influence of temporal envelope coding on pure-tone corrected speech comprehension thresholds, we analyzed ASSR for 4 kHz and 6 kHz carrier frequencies, and a modulation frequency of 116 Hz – the same frequency that was used as the fundamental frequency for the speech stimuli. We took into account that contributions of ASSR sources in response to lower (10 - 40 Hz) modulation rates are dominated by cortical components, while ASSR in response to higher (> 100 Hz) modulation rates are dominated by subcortical components (Engelien et al., 2000; Kuwada et al., 2002; Purcell et al., 2004; Lu et al., 2022). Testing ASSR for 116 Hz amplitude modulation therefore provides insight into the temporal processing of acoustic stimuli in subcortical, rather neocortical areas. The averaged ASSR amplitudes are shown as a function of age (Fig.8 A), and inspected for significance in groups with poor, standard, and good speech comprehension (orange, grey and blue dots) (Fig. 8 B).

**Figure 8.**
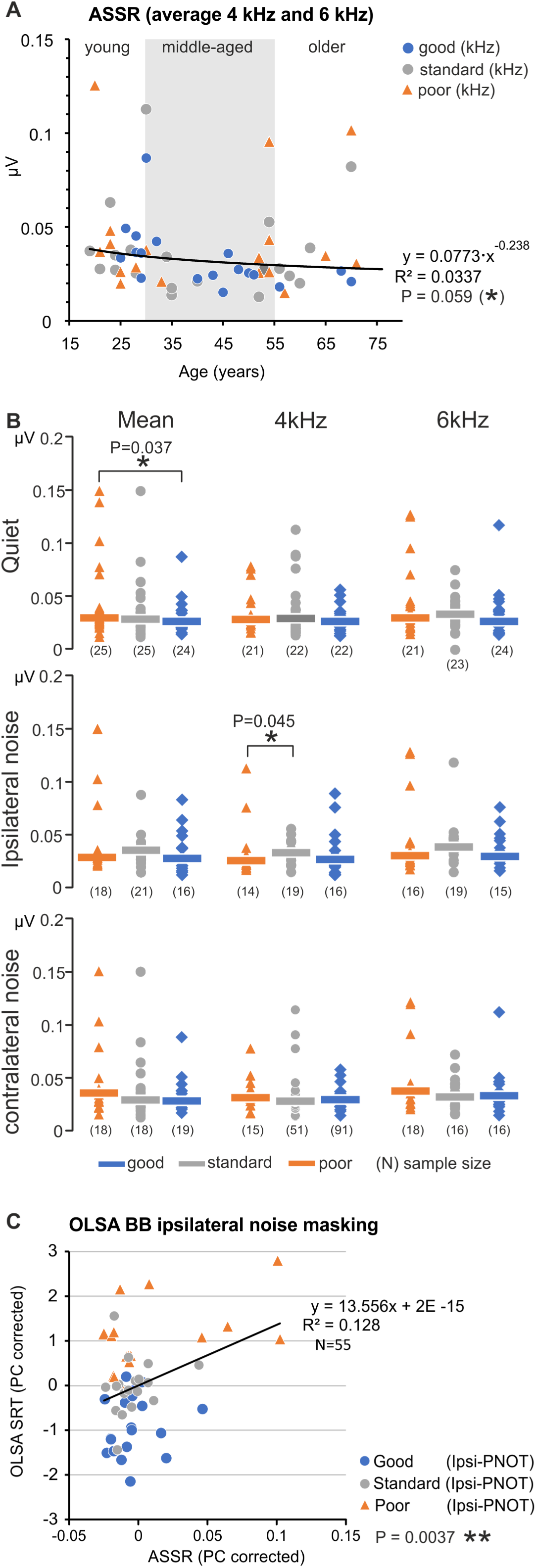
ASSR response amplitudes for 116 Hz amplitude modulated carriers at 4 kHz and 6 kHz. (**A**) Averaged (4 kHz and 6 kHz) responses (µV) over age (years) for particiant categorised by PNOT in quiet as good (blue,), standard (grey), and poor (orange) speech comprehension groups. The trendline (black) indicates smaller ASSR amplitude values with higher age, the correlation that just fails to reach statistical significance (corresponding R2 value and P-value for Pearson correlation coefficient). The shaded gray area marks the middle-aged group. (**B**) Individual ASSR amplitudes (symbols) and median (horizontal bar) for the respective speech comprehension groups categorized by PNOT in quiet (top row), ipsilateral noise masking (middle row), and contralateral noise masking (bottom row). Group differences significant for non-parametric statistical tests are marked by brackets and P-value in the respective panel (*, P<0.05 Mann-Whitney U test). (**C**) The regression of PCA-corrected speech reception threshold (OLSA SRT) and PCA-corrected ASSR response amplitude revealed a significant association (corresponding R2 value and P-value for Pearson correlation coefficient, ** P<0.01) between SRT and ASSR response amplitude with remarkably higher ASSR amplitudes in participants categorised as poor speech comprehension group for their SRT in ipsilateral noise masking (PNOT in ipsilateral noise). N=74 and 55 in (A) and (B), respectively. Selected ASSR response amplitudes for SNR>2.0 and µV<0.15, see Methods.

A slight negative correlation of lower amplitude with increasing age was noted for ASSR amplitudes at both 4 and 6 kHz carrier frequencies (shown for the average of 4 kHz and 6 kHz ASSR responses in Fig. 8 A). The four subjects who were outside the normal levels in the BDI and GDS tests (Fig. 1) were distributed within normal levels for all ASSR conditions (Fig. 8, black circled dots). The grouped ASSR amplitudes were found to be significantly different (larger) in the group with poor speech comprehension in quiet for the mean ASSR response (Fig. 8 B, top left panel, p=0.037) and in ipsilateral noise masking for 4 kHz (Fig. 8 B, middle panel, p=0.045). Interestingly, also in the *post hoc* linear-mixed model analysis after permutation, ASSR was the only electrophysiological measure that significantly explained a considerable amount (7%) of speech-in-noise comprehension (p = 0.012); this corresponded to 0.4 dB in the broadband condition (Tab. 4, Fig. 3). Therefore, correcting SRTs and ASSR amplitudes for PTT -related variance using the normalization by PCA revealed a significant association between ASSR amplitude and speech recognition threshold (Fig. 8 C, p=0.0037).

### Subjects with good and poor speech comprehension differ in cochlear-amplifier status

To determine whether good and poor speech-comprehension groups after normalization for PTT is accompanied by subtle threshold changes associated with cochlear-amplifier performance, we analyzed short-pulse DPOAE growth functions to determine the state of the cochlear amplifier with high accuracy (Zelle et al., 2017).

Extrapolated DPOAE thresholds based on semi-logarithmically scaled I/O functions have been shown to correlate nearly 1:1 with pure-tone threshold (Boege and Janssen, 2002; Gorga et al., 2003; Johnson et al., 2007; Zelle et al., 2017) when stimulus levels of both primaries are chosen according to a so-called scissors paradigm (Kummer et al., 1998). In contrast, DPOAE amplitudes as a stand-alone measure bear a more complicated, frequency- and level-dependent relationship to behavioral thresholds (Boege and Janssen, 2002; Gorga et al., 2003; Johnson et al., 2007; Zelle et al., 2017). The standard deviation of the residual of the correlation between L_EDPT_and pure-tone Békésy-tracking audiometry at f_2_ has been shown to range between 5 and 8 dB depending on frequency (Zelle et al., 2017). All of the above-mentioned studies used several criteria for I/O functions to be accepted, which effectively avoid hard-to-interpret DPOAE I/O functions that lead to large estimation errors at the cost of losing information. Of course, any inter-individual variation in synaptic and neural transmission as well as variation in performing the subjective threshold-detection task contributes to the prediction error. It is and has been understood that the measured DPOAE threshold is merely a predictor for the state of the cochlear amplifier, or, with other words, a predictor for the pre-neural input signal to the mechano-electrical transduction of the inner hair cells (Zelle et al., 2017).

When DPOAEs were analyzed in participants that were classified by good (Fig. 9A-D, blue) or poor (Fig. 9A-D, orange) speech-in-quiet recognition, four out of ten factors became significant, and one became a tendency. **(i)** The percentage of accepted estimations of p-DPOAE thresholds, L_EDPT_, were higher for participants with good speech recognition in comparison to participants with poor speech recognition for both the left (n = 60 ears; p = 0.039) and right (n = 60 ears; p = 0.039) ear (Fig. 9A). The difference in acceptance rate was assessed with a chi-squared test, using the two speech performance groups as the first dimension and above or below-average L_EDPT_ acceptance rate as a second dimension. **(ii)** The PTT was not different between groups (Fig. 9B). **(iii)** For the left ear, L_EDPT_ was significantly lower for the good performers (p = 0.012) (Fig. 9C). **(iv)** When L_EDPT_ were normalized for PTT, significantly lower cochlear threshold persisted in participants with good speech comprehension in comparison to those with poor speech comprehension in the left ear (p = 0.017) and remained different with a tendency (p = 0.084) for the right ear as well (Fig. 9D). The difference in normalized L_EDPT_-PTT between both groups was 2.84 and 3.14 dB for the right and left ear, respectively (Fig. 9D). Even when excluding the results for 8 kHz, higher L_EDPT_-PTT differences were observed for subjects with poor speech performance, showing again significance in the left and a tendency in the right ear.

**Figure 9.**
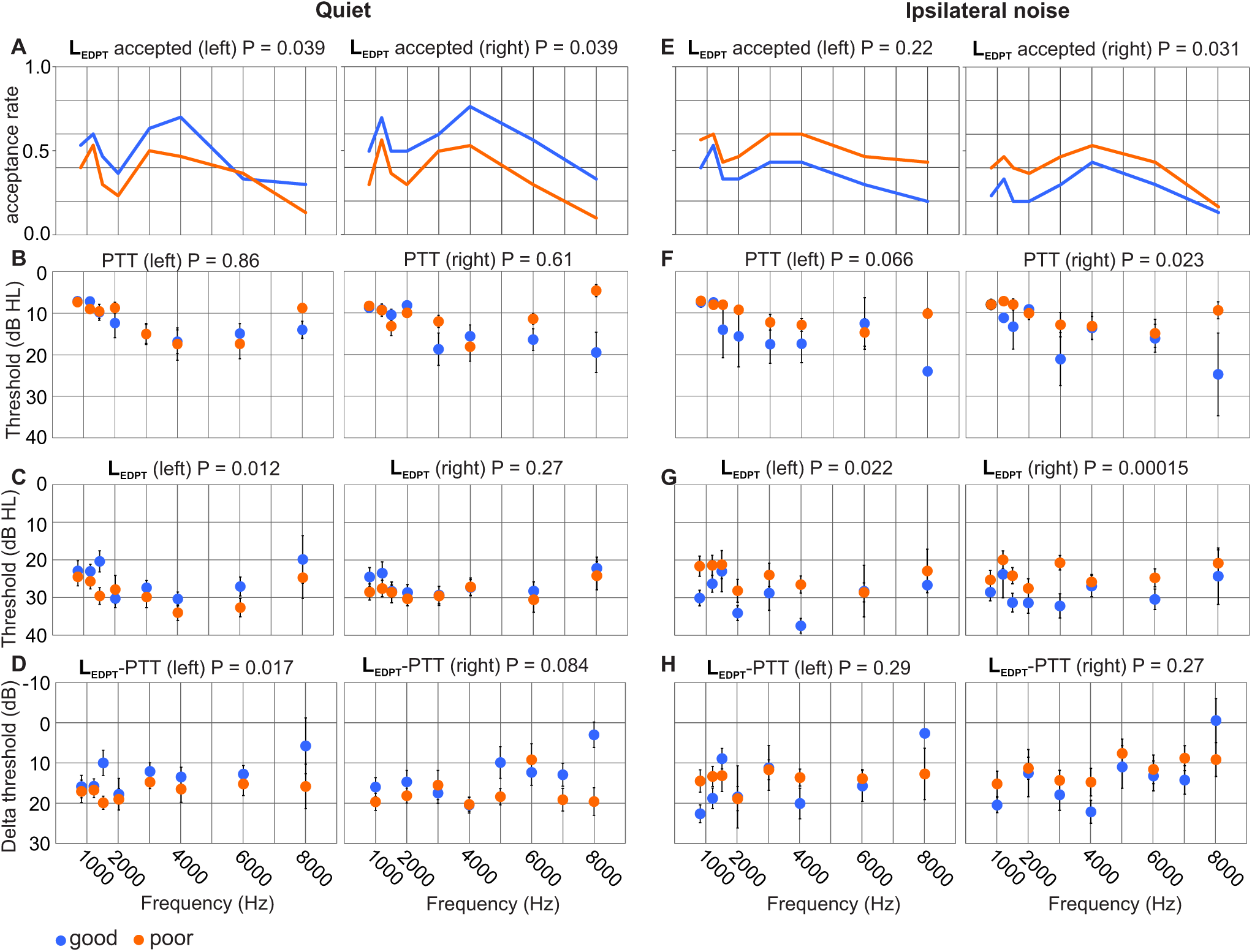
Estimated distortion-product thresholds (LEDPT) in relation to PTT in quiet (A-D) and ipsilateral noise (E-H). (A) LEDPT acceptance rates, (B) Pure tone thresholds (PTT), (C) LEDPT, (D) LEDPT-to-PTT difference for left and right ears are compared between participants with good (blue) and poor (orange) speech-in-quiet comprehension.. Participants with good speech-in-quiet performance show higher acceptance rates, equal PTT, inconclusive LEDPT, but a consistently 3 dB better threshold for PT-normalized LEDPT although on the right ear with only p = 0.084. E-H in ipsilateral noise (E) LEDPT acceptance rates, (F) PTT, (G) LEDPT, (H) LEDPT -to-PTT difference for left and right ears are compared between good (blue) and poor (orange) speech-in-noise comprehension performers. Participants with good speech-in-noise performance (blue) show reduced acceptance rates, reduced PTT and LEDPT, but no difference between PTT and LEDPT.

Thus, in a nutshell, semi-logarithmic DPOAE I/O functions, a measure which relates to cochlear amplification at near-threshold sound pressure levels (L_EDPT_), as well as a measure which is at least influenced by cochlear amplification at stimulus levels up to 55 dB SPL (acceptance rate), represents a stronger cochlear amplifier (lower L_EDPT_ values, higher acceptance rates) if a subject has good speech-in-quiet recognition or weaker amplifier if a subject has poor speech-in quiet recognition. It is, however, curious that the finding with respect to L_EDPT_ and the difference between L_EDPT_ and PTT obtained higher significance in the left ear, being the contralateral side to the ear in which speech recognition was assessed. If one disregards the lack of complete consistency, the conclusion would be that a stronger pre-neural input signal is an advantage for speech-in-quiet recognition for subjects with equal behavioral thresholds, and thus could point to an until now “hidden” influence of cochlear amplification in speech-in-quiet recognition.

When DPOAE were analyzed in participants that were classified by good (Fig. 9E-H, blue) and poor (Fig. 9 E-H, orange) speech-in-ipsilateral-noise recognition, five out of ten factors became significant, and one a tendency (only four measures for two ears shown in Fig. 9; we omitted the slope of the I/O functions that, with one exception, never became significant). In contrast to quiet conditions, the acceptance rate was significantly higher for poor performers in the right ear (p = 0.031) (Fig. 9E), the PT hearing threshold was lower for poor performers, with a tendency in the left ear (p = 0.066) and significant in the right ear (p = 0.023; Fig. 9F). Moreover, L_EDPT_ were significantly lower for poor performers in the left (p = 0.0022) and the right ear (p = 0.00015, Fig. 9G). In addition, the slope in the right ear was significantly steeper for poor performers (p = 0.041, not included in the figures).

Thus, measures of hearing sensitivity close to threshold, the PTT, the distortion-product threshold (L_EDPT_), and a measure which is at least influenced by cochlear amplification at levels up to 55 dB SPL (acceptance rate), represent stronger cochlear amplification (lower L_EDPT_ values, lower behavioral thresholds, higher acceptance rates) if a subject has poor speech-in-noise recognition.

This finding for the speech-in-ipsilateral-noise condition is more significant than the seemingly converse finding for speech-in-quiet, as the significance, especially for L_EDPT_, is high, and significance or at least a tendency appears consistently in five out of six conditions. This supports the finding that, in contrast to subjects with poor speech comprehension in quiet, those with poor speech comprehension in noise do not have low but rather higher cochlear amplification performance.

When participants were classified by speech recognition in contralateral noise, one factor became significant and one became a tendency. For the left ear, the slope of the I/O function was significantly steeper for the good performers (p = 0.039) and the difference between L_EDPT_ and PTT was found to be higher for the bad performers, with a tendency in the right ear (p = 0.084). Thus, for speech recognition in contralateral noise, there is no obvious rule relating any of these measures to speech recognition advantage or disadvantage, except the solitary finding of I/O function slope in the ear contralateral to the speech recognition testing (not shown in figures). Thus, for participants with poor speech comprehension in ipsilateral noise, a better (lower) threshold may be characteristic, which was caused as a “hidden” compensatory reaction to the same damage that may also contribute to a poor speech comprehension quality in contralateral noise.

Finally, we tested whether there were contributions to the total variance of speech comprehension performance based in each of the three differently filtered versions of OLSA. We found that DPOAE I/O function acceptance rate (Fig. 9A) as well as the difference between L_EDPT_ and PTT (Fig. 9D) survived the most restrictive *post hoc* linear mixed model analysis after permutation (p = 0.001 - 0.033), explaining 2.0 - 8.3% of the variance, or 0.8 dB and 3.2 - 4.8 dB of SRT variation in the broadband and high-pass condition, respectively. Under ipsilateral noise, the acceptance rate of L_EDPT_ measurements (Fig. 9A) was significant to explain variance of SRT in broadband and high-pass condition and almost significant (p = 0.051) in the low-pass condition, accounting for 3.1 - 5.5% of the variance, but only 0.3 - 1.0 dB in SRT. Here, the PT corrected L_EDPT_ thresholds explained up to 7% (0.3 dB in SRT) of the variance of the OLSA, but only in the low-pass condition (Fig. 3, Tab. 4).

### Participants with good and poor speech comprehension differ in response latencies of auditory nerve fibers

Weaker cochlear amplification at sound pressure levels near threshold, as here observed in individuals with poor speech comprehension in quiet, may influence the dynamic range of inner hair cell excitation in quiet conditions. To measure this, we analyzed the amplitudes of suprathreshold ABR waves, which are defined through the precise discharge rate of individual auditory fibers (Buran et al., 2010), in particular the precision with which auditory fibers fire synchronously at onset (Johnson and Kiang, 1976). Synchronous firing rate at onset of the stimulus is a feature that critically dependent upon the sensitivity of high spontaneous firing rate (high-SR) low threshold auditory fibers (Rhode and Smith, 1986; Meddis, 2006; Heil et al., 2008; Zohar et al., 2011).

We thus analyzed differences in the onset of brainstem responses, comparing participants with good and poor speech comprehension based on PNOT. Amplitudes and latencies of suprathreshold ABR responses of ABR wave I (generated by the auditory nerve) (Portmann et al., 1980), ABR wave III (generated by the superior olivary complex (**SOC**) and lateral lemniscus) (Melcher et al., 1996), and ABR wave V and VI (generated by the inferior colliculus (**IC**) (Møller et al., 1994) and the medial geniculate body (**MGB**) (Hashimoto et al., 1981), were measured by acoustic stimulation at 80 dB SPL rms in young, (Fig. 10A, solid line, circle), middle-aged (Fig. 10A, dotted line, triangle) and older participants (Fig. 10A, dotted line, square). ABR wave I was significantly lower in the middle-aged and older participants than in the young participants (Fig. 10A, young: 0.203 µV, n = 27; middle-aged: 0.111 µV; n = 27; older 0.119 µV; n = 19; p = 0.00029), indicating an age-dependent cochlear synaptopathy. However, this lower input was somewhat compensated by ABR wave VI in the middle-aged group (Fig. 10A, young: 0.355 µV, n = 38; older: 0.298 µV, n = 40; p = 0.041) but was still significantly delayed in the older group (Fig. 10A, wave III, youg: 3.67 ms, n = 29; middle-aged: 3.77 ms; n = 31; older: 3.86 ms, n = 24; p = 0.0095; wave V, young: 5.55 ms, n = 29; middle-aged: 5.66 ms, n = 32; older: 5.76 ms, n = 25; p= 0.018). The lower-amplitude and delayed ABR wave VI in the older group was not linked to PTT differences in the PTA-4 frequency range (Fig. 10B) or the PTA-HF frequency range (not shown), but rather to lower ABR wave V and VI thresholds (Fig. 10C) and significantly delayed latency differences at EHFs (Fig. 10C). Analyzing the ABR parameters for PNOT groups (Fig. 10D), we observed that the amplitude of wave I differed by 0.0148 +/- 0.0129 µV in participants with poor speech comprehension in comparison to those with good speech comprehension, which limits the detection threshold for input amplitude differences to 5 dB when assuming the ABR wave I growth with respect to volume as described by (Eggermont and Odenthal, 1974). Moreover, we found a significantly smaller ABR wave II amplitude in participants with poor speech comprehension in comparison to those with good speech comprehension (Fig. 10D, good: 0.0767 µV, n = 24; standard: 0.0905 µV, n = 22; poor: 0.0458 µV, n = 16; p = 0.0458), while no difference in the ABR wave III, V, or VI amplitude between participants with good and poor speech comprehension was observed (Fig. 10D).

**Figure 10.**
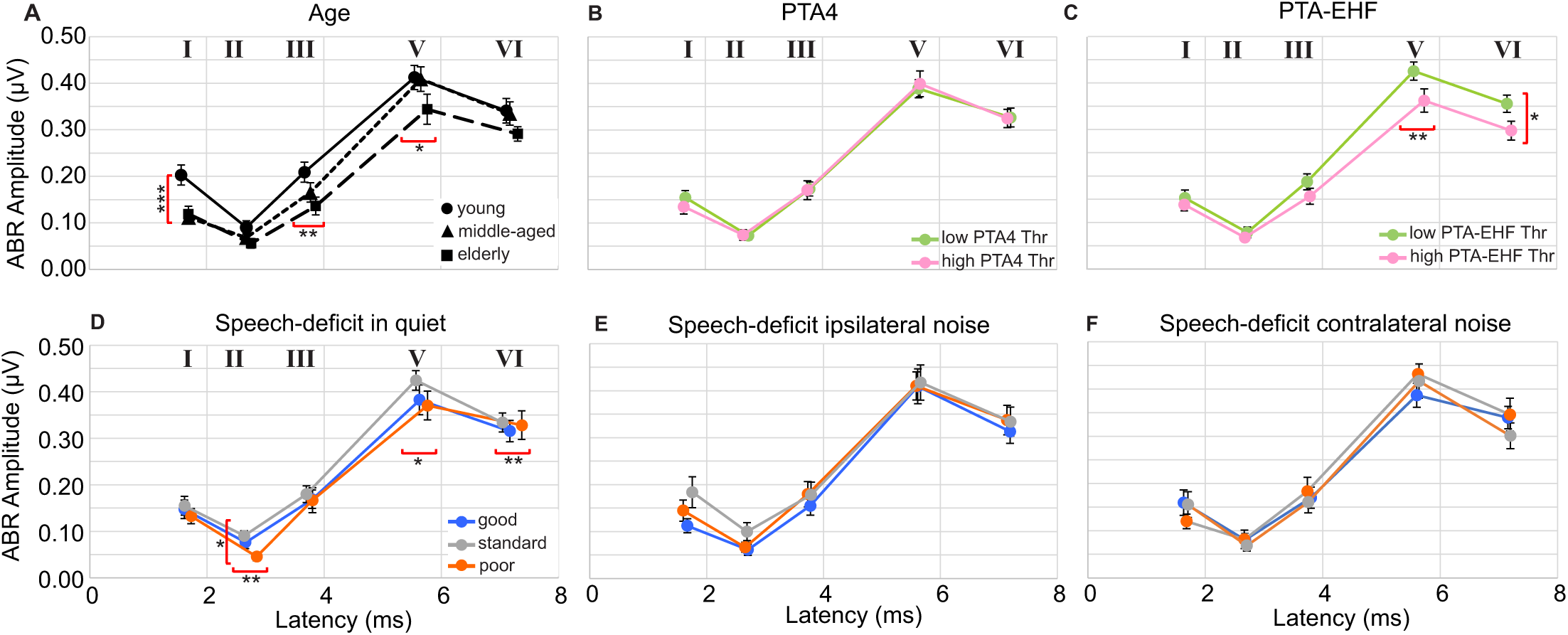
ABR as a function of age, pure-tone averages, and speech comprehension. (A) ABR wave amplitude as a function of age. Circles: young group; triangles: middle-aged group; squares, older group. (B) and (C) display ABR waves as function of low (pink) versus high (green) threshold of PTA4 (B) and PTA-EHF (C). For individuals with matched PTA-thresholds for good (blue), standard (grey), or poor (orange) speech comprehension in quiet (D), ipsilateral noise (E), and contralateral noise (F), suprathreshold ABR wave amplitudes and latencies were determined. For speech comprehension in quiet (D), significant shifts in latency in the group with good speech comprehension in comparison to the group with poor speech comprehension was observed (ABR wave I: n = 29, 27, 24, p = 0.218242; ABR wave II: n = 24, 22, 16, p = 0.007707; ABR wave III: n = 30, 28, 26, p = 0.182784; ABR wave V: n = 30,28, 28, p = 0.026617; ABR wave VI: n = 27, 27, 24, p = 0.001055).

Interestingly when testing the predictive ability of ABR waves for variance of speech reception thresholds in quiet (Fig. 3, Tab. 4) we found that the ABR wave I amplitude was negatively correlated with the OLSA-BB and OLSA-LP OLSA explaining 2.5 and 2.8% of the variance, respectively, while no effects could be observed under ipsilateral noise (Fig. 3, Tab. 4).

In contrast, significant latency shifts were observed in participants with poor speech comprehension in comparison to those with good or standard speech comprehension, as shown for wave II (Fig. 10D, good: 2.66 ms, n = 24; standard: 2.63 ms, n = 22; poor: 2.85 ms, n = 16; p = 0.00771), wave V (Fig. 10D, good: 5.63 ms, n = 30; standard: 5.56 ms, n = 28; poor: 5.76 ms, n = 28; p = 0.027), and wave VI (Fig 10D, good: 7.17 ms, n = 27; standard: 7.04 ms, n = 27; poor: 7.37 ms, n = 24; p = 0.0011).

Surprisingly, the latency difference in the good and poor PNOT groups in quiet did not differ by age (Tab. 5) or by PTA-EHF (Tab. 5). This indicates that the delayed ABR wave latency in groups with poor speech comprehension in quiet (Fig. 10D) is not due to age difference or to elevated PTA-EHF levels.

When PNOT groups were classified for good and poor speech comprehension in ipsilateral noise (Fig. 10E) or contralateral noise (Fig. 10F), neither ABR wave II amplitude nor absolute latency differed between the groups. These results also did not differ by age or PT-EHF when corrected for missing data (Tab. 5).

In addition, we observed that ABR latencies contribute to SRTs and found that ABR wave I and V survived the most restrictive *post hoc* linear mixed model analysis after permutation (p = 0.004 - 0.030), and significantly explained 2.2% and 2.2% of OLSA-BB variance and 4.6% and 3.2% of OLSA-LP variation in quiet, however, only corresponding to 0.8 - 1.3 dB of SRT variation. ABR wave VI latency explained 4.2% of the OLSA-HP in quiet (Fig 3, Tab. 4). No significant contributions of the ABR wave could be observed under ipsilateral noise for any of the conditions. Among the four subjects who were outside the normal levels in the BDI and GDS tests (Fig. 1), two belonged to the group with poor speech comprehension in quiet group, exhibiting prolonged latency, and two belonged to the group with poor comprehension in ipsilateral noise (Fig. 10).

**We conclude (i)** that compensatory central neural gain following a cochlear synaptopathy (i.e., the disproportional increase of the ABR wave V amplitude to the ABR wave I amplitude (Fig. 10A) is dependent upon PTA-EHF hearing (Fig. 10C). **(ii)** The compensatory central neural gain is not related to speech comprehension when OLSA thresholds are normalized for PTT. **(iii)** Poor speech comprehension in quiet (Fig 10D) not in noise Fig. 10E, F) is linked with delayed sound responses and slightly lower ABR wave II amplitude.

### Participants with good and poor speech comprehension differ in phoneme discrimination

We have thus far identified “hidden” performance differences of the cochlear amplifier and associated differences in latency of ABR waves between participants exhibiting good or poor speech comprehension in quiet or ipsilateral noise, despite no obvious changes in PTT. We aimed to investigate a relationship between these events by measuring phoneme discrimination ability in the different groups. Language comprehension is dependent upon correct discrimination of vowels (Won et al., 2016) and consonants (Hornickel et al., 2009) that in turn require precise temporal fine-structure coding (below the human phase locking limit; < 1500 Hz) and temporal envelope coding (above the phase locking limit; > 1500 Hz) (Weiss and Rose, 1988; Verschooten et al., 2019). We specifically aimed to test the groups with good and poor speech comprehension for differences in temporal sound coding strategies using a three-alternative forced choice psychoacoustic task to discriminate four different pairs of phonemes: /o/-/u/, /du/-/bu/, /i/-/y/, and /di/-/bi/. The /o/-/u/, /du/-/bu/ phoneme pairs had formant contrasts below the human PLL at 1500 Hz – enabling temporal fine structure coding – while the /i/-/y/ and /di/-/bi/ phoneme pairs had the contrasts above the PLL –requiring temporal envelope coding (Tab.1). The consonant-based phoneme contrasts were thus assigned to the two different processing mechanisms, /du/-/bu/ to temporal fine structure coding and /di-bi/ to temporal envelope coding.

All phoneme pairs were presented in randomized blocks to the right ear in quiet, ipsilateral noise, and contralateral noise. For all tested phoneme pairs, two grades of difficulties were chosen depending on the size of the physical contrast (here labeled as “difficult” and “easy”). When the discrimination ability in percent is plotted against age for phoneme pair discrimination in quiet, ipsilateral noise, and contralateral noise conditions, a weak correlation was found in the quiet condition for /di/-/bi/ and in the ipsilateral noise condition for /du/-/bu/ (data not shown).

The ability to discriminate between the indicated phoneme pairs in quiet, ipsilateral noise, and contralateral noise was plotted against the PNOTs obtained from the corresponding groups (Fig. 11). Both, easy and difficult discrimination conditions were averaged. Tab. 7 provides statistics on behavioral accuracy.

**Figure 11.**
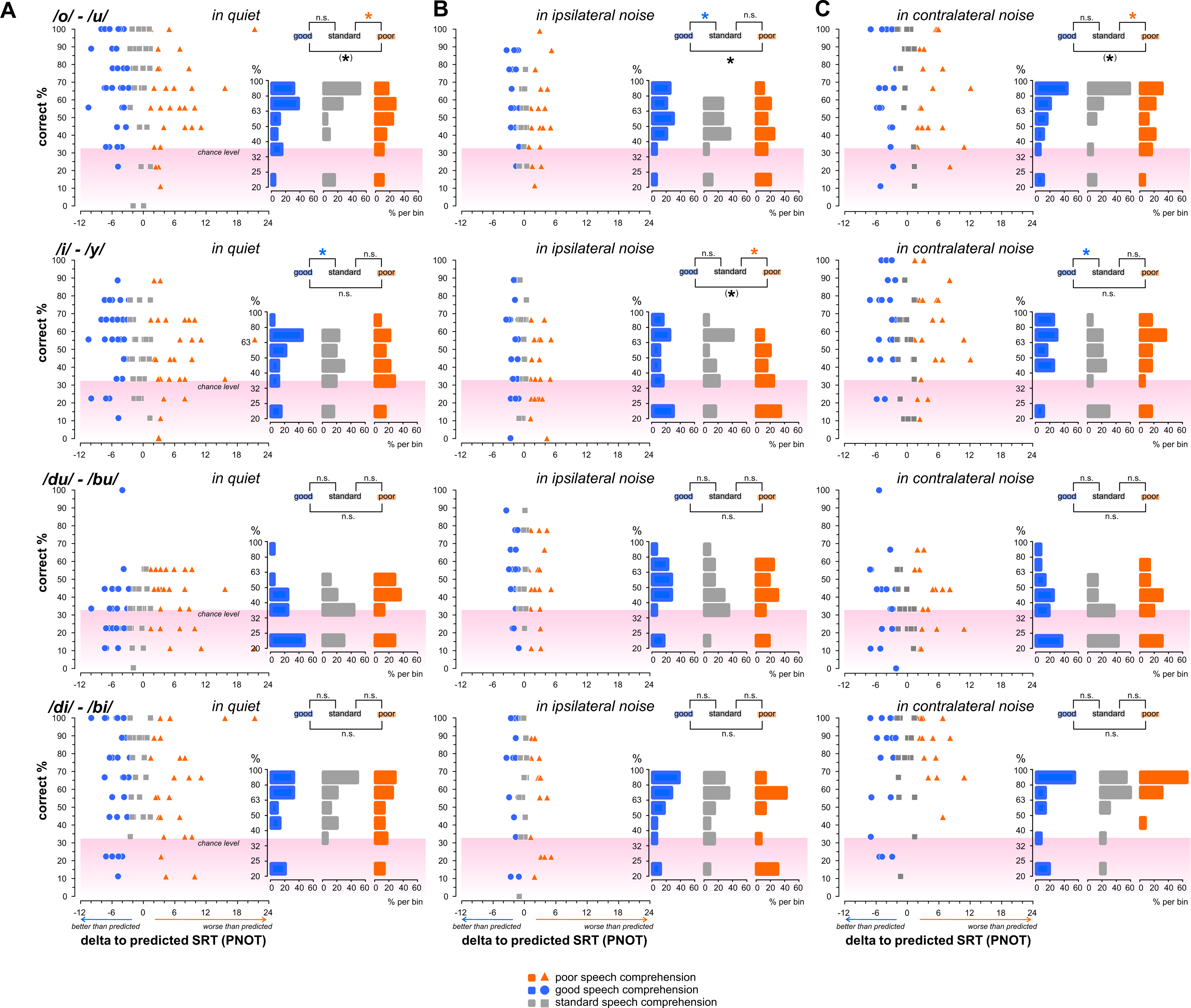
Syllable discrimination scores for 4 pairs of phonemes in relation to speech comprehension. Syllable discrimination scores for 4 pairs of phonemes (/o/-/u/, /i/-/y/, /du/-/bu/, /di/-/bi/) segregated for participants with poor (orange), good (blue) and standard (grey) speech comprehension selected by PNOT in quiet (A), ipsilateral noise (B) and contralateral noise (C). Each subplot consists of a scattergram (left) with perceptual performance [% correct] as a function of PNOT (x-axis). The right side of each subplot consists of a frequency distribution along the y-axis, segregated for the three speech comprehension groups in the same colors as in the scattergram on the left. The counts are converted into percent from counts per bin. Finally, there is a graphical representation of the significance assessed by Mann-Whitney-U-tests (Table 6), providing significant differences as asterisks with a color code reflecting the three groups.

**Table 6.** P-values for syllable discrimination scores for different speech performers. Test of significance by 1-sided Mann-Whitney U2 test for syllable discrimination scores between good, standard, and poor speech performers, classified by PNOT. Bold values, p<=0.05. Difficult, small spectral and temporal envelope contrast. Easy, large spectral and temporal envelope contrast. Poor, Standard, Good, group category from speech recognition threshold in quiet, ipsilateral, and contralateral noise (PNOT).

| listening condition |  | quiet |  |  | ipsilateral noise masking |  |  | contralateral noise masking |  |  |
| --- | --- | --- | --- | --- | --- | --- | --- | --- | --- | --- |
| group | comparison | poor | poor | good | poor | poor | good | poor | poor | good |
|  |  | vs. | vs. | vs. | vs. | vs. | vs. | vs. | vs. | vs. |
|  |  | good | standard | standard | good | standard | standard | good | standard | standard |
| /o/ - /u/ | (difficult) | 0.059 | 0.011 | 0.159 | 0.032 | 0.337 | 0.035 | 0.125 | 0.047 | 0.386 |
| /i/ - /y/ | (difficult) | 0.496 | 0.444 | 0.041 | 0.081 | 0.031 | 0.433 | 0.316 | (0.025) | 0.008 |
| /du/ - /bu/ | (difficult) | (0.0314) | (0.0584) | 0.334 | 0.248 | 0.448 | 0.233 | 0.378 | 0.121 | 0.245 |
| /di/ - /bi/ | (difficult) | 0.291 | 0.102 | 0.264 | 0.042 | 0.187 | 0.189 | 0.0951 | (0.0869) | 0.480 |
| /o/ - /u/ | (easy) | 0.007 | 0.001 | 0.264 | 0.212 | 0.337 | 0.125 | 0.031 | 0.013 | 0.195 |
| /i/ - /y/ | (easy) | 0.014 | 0.084 | 0.236 | 0.129 | 0.248 | 0.356 | 0.337 | 0.161 | 0.284 |
| /du/ - /bu/ | (easy) | 0.076 | 0.284 | 0.271 | 0.179 | 0.061 | 0.245 | 0.239 | 0.221 | 0.464 |
| /di/ - /bi/ | (easy) | 0.264 | 0.176 | 0.378 | 0.284 | 0.326 | 0.492 | 0.176 | 0.302 | 0.323 |

In general, performance of all participants was better for the discrimination of /di/-/bi/ than of /du/-/bu/. Thus, the phoneme /du/-/bu/ (Fig 11, A,B,C /du-bu/) showed the slimmest variation of behavioral results across the cohort, with performance exceeding the 66^th^ percentile only by a single participant (CS083), regardless of age, and on average 29.4% to 58.9% of the participants responding below or at the 33th percentile mark (= chance level), dependent on the noise condition. The highest percentage of correct behavioral responses was achieved for discrimination of /o/-/u/ and /di/-/bi/, less for /i/-/y/ (Fig. 11A,B,C).

All phoneme pairs were presented in randomized blocks only on the right ear in quiet, ipsi- and contralateral noise (see methods). For all tested phoneme pairs, two grades of difficulties dependent on the size of the physical contrast were chosen (here labeled as “difficult” and “easy”). When the discrimination ability in percent is plotted against age for phoneme pair discrimination in each listening condition, a weak correlation was found in the quiet condition for /di/-/bi/ and in the ipsilateral noise condition for /du/-/bu/ (data not shown).

The ability to discriminate between the indicated phoneme pairs in quiet, ipsi and contralateral noise was plotted against the PNOTs (Fig. 11) obtained from the correspondent groups categorized in quiet, ipsi or contralateral noise conditions. The easy and difficult discrimination conditions were averaged for the individual condition. Tab. 7 provides statistics on behavioral accuracy.

In general, performance of all participants was better for the discrimination of /di/-/bi/ than for /du/-/bu/. Thus, the phoneme /du/-/bu/ (Fig 11, A,B,C, /du/-/bu/) showed the least variation across the cohort. Only a single participant’s performance exceeded the 66^th^ percentile (CS083), and on average, regardless of age, 29.4% to 58.9% of the participants responded below or at the 33th percentile mark (i.e., chance level), depending on the noise condition. The highest percentage of correct responses was achieved for discrimination of /o/-/u/ and /di/-/bi/ (Fig. 11A,B,C).

The most prominent difference between participants with good or poor PNOT was in the differentiation of /o/-/u/, with formant contrasts below the PLL, as shown for quiet (Fig. 11A), ipsi-noise (Fig. 11B) or contralateral conditions (Fig. 11C). When PNOT categorization in quiet (Fig. 11A) or contralateral noise (Fig. 11C) was analyzed for phoneme discrimination, it became evident that the /o/-/u/ discrimination performance in participants with poor speech comprehension in quiet was worse than in the group with standard speech comprehension (Fig. 11A,C). Even under easy conditions, in which the two stimuli had large spectral differences, groups with poor speech comprehension in quiet performed worse in comparison to those with good or standard speech comprehension in quiet or contralateral noise (Tab. 6). On the other hand, participants with good speech comprehension (categorized from PNOT) in quiet (Fig. 11A, /i/-/y/) and contralateral noise (Fig. 11C, /i/-/y/) were significantly better in their discrimination of /i/-/y/ in comparison to participants with standard speech discrimination ability.

Under ipsilateral noise conditions, participants with good speech comprehension (categorized from PNOT) were better able to discriminate between /o/-/u/ than participants with standard speech comprehension. However, groups with poor speech comprehension were worse at discrimination between /i/-/y/ in comparison to those with standard speech comprehension (Fig. 11B, /i/-/y/) for both easy and difficult discrimination conditions in ipsilateral noise (Tab. 6).

The discrimination of /du/-/bu/ was not different between participants with good and poor speech comprehension (categorized from PNOT) in quiet, ipsilateral noise, or contralateral noise (Fig.11 A,B,C /du/-bu/), likely due to the fact that the performance rate among all participants almost never exceeded 60%.

Further, groups with good, standard, and poor speech comprehension (categorized from PNOT) did not differ in their discrimination ability between /di/-/bi/ in any of the listening conditions (Fig. 11 A,B,C /di/-/bi/), likely due to the fact that the performance rate among all participants almost always exceeded 90%. Important to say in this context that the numbers shown in Fig. 11 and Tab. 6 are not corrected for multiple comparisons and these effects are likely limited by saturation effects in task performance resulting in a non-linear dependence which our linear model of variance analysis cannot capture.

Thus, limits in exceeding dimensions for statistical analysis, allowed the variance analysis for phoneme discrimination only as linear dependency between good and poor PNOT, but not for the non-linear contribution of phoneme discrimination deficits in participants with good against standard or poor against standard speech comprehension. Different from results obtained from non-linear dependence (Fig. 11, Tab. 6), the linear dependencies for phoneme contrasts in good and poor PNOT resulted in significant explanation of variance in quiet for /di/-/bi/ easy 2.3%, /o/-/u/ difficult 2% and / o/-/u/ easy 2% and in ipsilateral noise for /i/-/y/ difficult contributing to 4.1% of variance (Tab. 4).

**Overall** our findings show that good and poor speech comprehension in quiet (and contralateral noise) differs from good and poor speech comprehension in ipsi-lateral noise in their discrimination ability of formant contrasts below PLL (requiring temporal fine structure coding) and above PLL (requiring temporal envelope coding) (Fig. 12).

**Figure 12.**
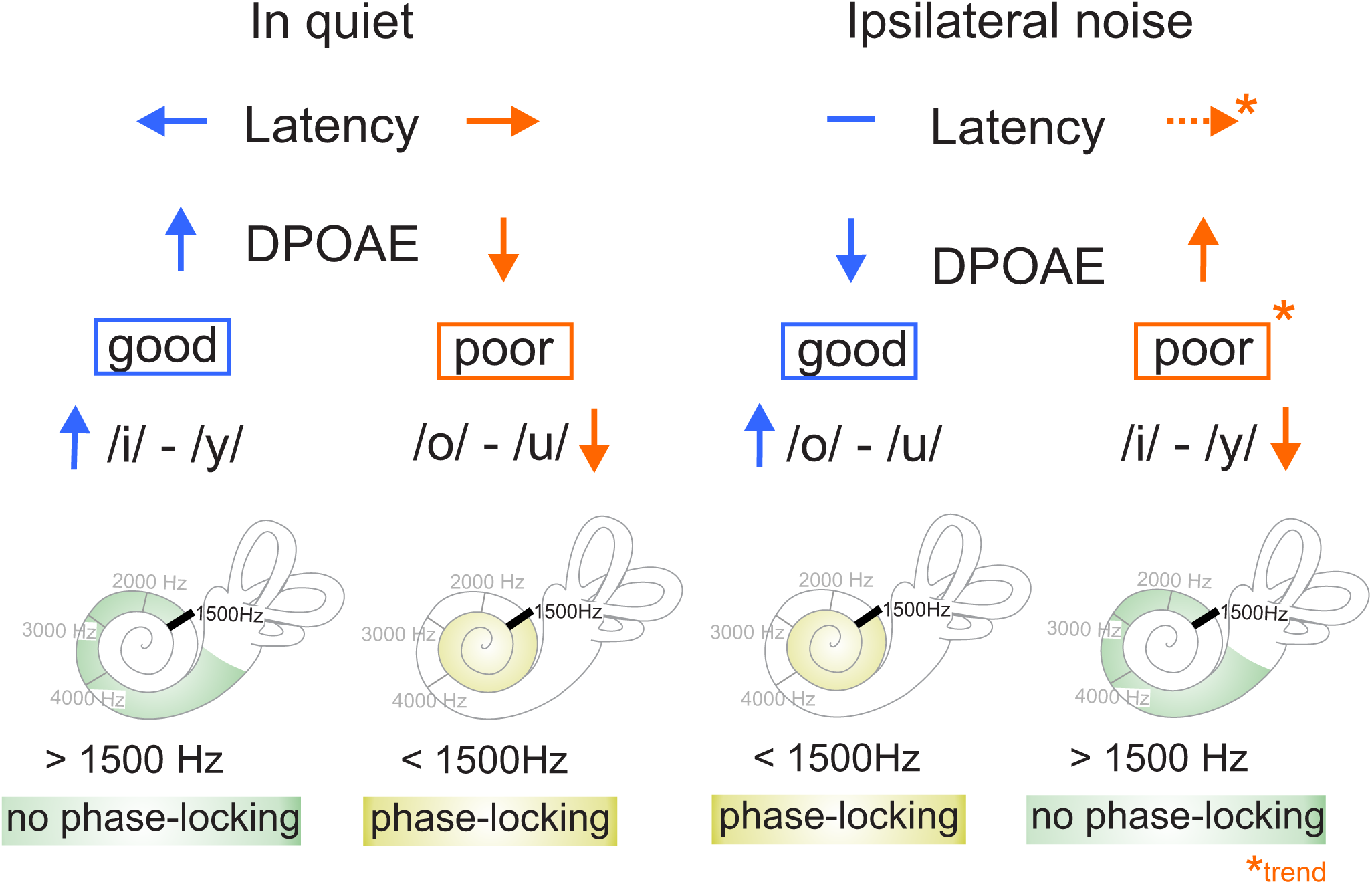
Graphical abstract: Good and poor speech comprehension in quiet differs from good and poor speech comprehension in ipsi-lateral noise in there discrimination ability of formant contrasts below PLL (requiring temporal fine structure coding) and above PLL (requiring temporal envelope coding).In quiet (or contra-lateral noise), poor speech comprehension is associated with poor discrimination below the phase locking limit (e.g. for /o-u/), while good speech comprehension is associated with good discrimination above the phase locking limit (e.g. for /i/-/y/). In ipsi noise, poor speech comprehension is associated with lower performance for discriminating phoneme pairs with formant contrasts above the phase-locking limit (/i-y/>1500 Hz), while good speech comprehension is associated with good discrimination of formants below phase locking limit (/o-u/ < 1500 Hz) (Fig. 11, in noise).

**In quiet** (or contra-lateral noise), poor speech comprehension is associated with poor discrimination of phoneme pairs with formant contrasts below the PLL (/o/-/u/), while good speech comprehension is associated with better discrimination of phoneme pairs with formant contrasts above the PLL (/i/-/y/) (Fig. 12, in quiet).

**In ipsilateral noise**, poor speech comprehension is associated with poorer discrimination of phoneme pairs with formant contrasts above the PLL (/i/-/y/), while good speech comprehension is associated with good discrimination of phoneme pairs with formant contrasts below phase locking limit (/o/-/u/) (Fig. 12, in noise).

The differentiation of consonant-based phoneme contrasts that require temporal fine structure coding (i.e., /du/-/bu/), were too difficult for both good and poor PNOT groups, while the phone contrasts that require temporal envelope coding (i.e., /di/-/bi/) were too easy for both good and poor PNOT groups. Neither of these stimulus pairs therefore resulted in any group differences. Thus, the dynamic range of the phoneme task as implemented here was insufficient for differentiating the influence of PNOT for /du/-/bu/ and /di/-/bi/ for the different speech-coding mechanisms (temporal fine structure versus temporal envelope coding).

## DISCUSSION

The present study investigated contributing factors of SRT for young, middle-aged, and old participants with mostly normal hearing or mild hearing loss. We found PTT to be the dominant factor for SRT, explaining approximately half of the variance in quiet and in noise. The remaining variance was explained by a new measure independent of age that removed PTT influence from SRT (“PNOT”). We show a previously undescribed differential influence of the cochlear amplifier on speech comprehension in quiet and noise through ANF adaptation characteristics at the onset of the stimulus. This mechanism, along with the ABR wave I amplitude, latency, and ASSR explained part of the SRT variance in PNOT with possible differential impact on temporal fine structure or envelope coding.

### PTT and SRT show age dependent differences

In line with previous studies (Fullgrabe et al., 2014; Hunter et al., 2020; Wu and Liberman, 2022), we observed slight hearing loss in lower frequency ranges (PTA4 and PTA-LF) and prominent hearing loss at high and EHF (PTA-HF and PTA-EHF) over age (Fig.4). When subjects were grouped into three categories based on their PNOT (poor, standard, and good), there was a better correlation between these categories and the subjects’ self-reported evaluation of their comprehension ability when listening to speech in quiet than that when subjects were categorized by age (Figure 7). Although this correlation was only a statistical tendency it strengthen previous findings of no correlation between self-reported speech comprehension ability and age (Shehabi et al., 2022).

### Speech comprehension in quiet

Although SRTs in quiet strongly depends on PTT, it nevertheless leaves 38.7% of variance of OLSA-BB unexplained which corresponds to a SD of 3.7 dB in SRT. The variance left unexplained could partially be explained by other measures.

**First**, the difference between L_EDPT_ and PTT was a significant factor that explained 2% and 8.3% of the variance in OLSA-BB and OLSA-HP, respectively. A plausible reason for this effect is that L_EDPT_, which is in general closely related to PTT, is not subject to adaptation of the ANF firing rate (Kiang et al., 1966) or to adaptation caused by the medial olivocochlear reflex since its time constant (Kim et al., 2001; Bassim et al., 2003) is well above the DPOAE stimulus pulse widths used here. On the other hand, the PTT as implemented in clinical audiometry is effectively integrating over ∼500 ms and thus reflects the adapted state of nerve firing (Goutman, 2017). This would explain the fact that after removal of PTT influence by PNOT, the difference between both metrics remains a significant factor explaining variance, and it thus would reflect different nerve adaptation characteristics at the onset of the stimulus. In mammalian inner hair cells nerve adaptation at the onset is associated with vesicle depletion characteristic linked to synaptic fatigue or desensitization kinetic of postsynaptic receptors (Moser and Beutner, 2000; Goutman, 2017; Peterson et al., 2018; Willmore and King, 2023). Thus, a larger or smaller L_EDPT_-PTT differences that correlate with better or worse speech-in-quiet comprehension (Fig. 9A,D), would reflect stronger or lower firing-rate adaptation, linked to less or more synaptic fatigue at inner hair cells, being a plausible mechanism for differences in detection of signal-onset features.

**Second,** ABR wave I amplitude and latency differences contributed to PTT-corrected SRT, independent of age. It is important to mention that we found in this study, in line with previous work (Parthasarathy et al., 2019; Bramhall, 2021; Vasilkov et al., 2021), what has been called age-dependent synaptopathy: ABR wave I amplitudes were reduced in middle-aged and older subjects (Fig. 10A) but could not be compensated in the older. This lack of compensation was linked to a considerable loss of PTA-EHF (Fig. 10C), which may indicate that good EHF thresholds contribute to speech comprehension (Motlagh Zadeh et al., 2019).

However, what we might call functional synaptopathy, i.e., ABR wave changes statistically explaining part of the hidden hearing loss (Fig. 3, Tab.4), was not age-dependent. This functional synaptopathy was evident as an ABR wave I amplitude reduction and latency shift in participants with poor PNOT in quiet (Fig. 10D), which explained 2.8% – 1.0 dB – of OLSA-LP (ABR wave I) and 4.6% – 1.3 dB – of OLSA-LP (ABR wave I latency) when SRT was corrected for PTT (Tab. 4). Both reduced ABR wave I amplitude and prolonged latency are best explained by poor high-SR ANF performance, as this fiber class defines amplitude size through its synchronization of discharge rate (Johnson, 1980) and defines the shortest latency responses through its firing rate at characteristic frequency (Meddis, 2006; Heil et al., 2008). In addition to envelope coding, high-SR ANF play a decisive role for perceptional threshold in the phase-locking range (Bourien et al., 2014; Huet et al., 2018), that must be seen in connection with ABR wave I reduction and wave I-V delay in the group with poor speech comprehension in quiet (Fig. 10A) and worse speech coding below the PLL limit (Fig. 11A, /o/-/u/, Fig. 12). On the other hand, we found significantly better discrimination of phonemes with formant contrasts above PLL in subjects with good speech discrimination, which may be attributed to altered L_EDPT_ vs. PTT difference, explaining the variance in OLSA-HP (Fig. 11 A, /i/-/y/, Fig. 12). However, this was not entirely consistent, as significant effects were only observed in the easy condition for formants above PLL (/di/-/bi/) and for formants below PLL (/o/-/u/) for OLSA-BB, but not for OLSA-HP and OLSA-LP. Further investigation is obviously required to elucidate this aspect.

### Speech comprehension in ipsi-noise

SRT in ipsilateral noise also depends on PTT but leaves more of the variance unexplained (i.e., 51.4%, corresponding to a SD of 1.0 dB). We identified two more major contributors to the remaining variance. (1) L_EDPT_-PTT and the acceptance rate of DPOAE I/O functions (Fig. 9, Tab. 4) explained 7.0% and 5.5% of the remaining variance of SRT in the OLSA-LP condition, respectively, corresponding to 0.3 dB in SRT for both measures, and explained 0.7 and 3.1% in the OLSA-HP condition, respectively, corresponding to 0.5 and 1.1 dB in SRT. The sign of the variance of L_EDPT_-PTT and the acceptance rate swapped in comparison to the quiet condition (Fig. 9, Tab. 4), indicating that in ipsilateral noise, the larger DPOAE I/O function acceptance rate and larger L_EDPT_-PTT are linked to poorer speech comprehension (Fig. 9B). This finding might be explained by compression of the cochlear input signal to the neural system. For the speech-in-noise test, temporal information within only a narrow dynamic level range is used. If then the DPOAE I/O function acceptance rate is comparatively worse, the levels above which basilar-membrane compression basically ends and the growth behavior approaches linear dependency, would start at lower levels. This may be an advantage because a larger part of the dynamic level range used in the test would be almost linear, yielding uncompromised modulation contrast of the speech signal. If this contributes to hidden hearing loss, we here have encountered a well-known phenomenon which has historically been termed recruitment (Denes and Naunton, 1950) and is known to correlate to DPOAE I/O functions (Rasetshwane et al., 2013). As found in previous studies (Boettcher et al., 2001; Rumschlag and Razak, 2021) ASSR amplitude declined with age, although only showing a statistical tendency. We observed a higher ASSR amplitude in poor PNOT in ipsi-noise (Fig 8C) explaining 7% of the variance of speech-in-noise comprehension in the OLSA-BB condition, corresponding to 0.4 dB in SRT (Tab. 4). ASSR growth function is known to correlate well with loudness (Menard et al., 2008; Oxenham, 2018; Lu et al., 2022), and our data appear to be driven by a subgroup of poor performers who show extraordinarily high PC-corrected ASSR amplitudes (Fig. 8) and significant increase in UCL (Tab 4). This indicates that, possibly, part of the poor-performing group exhibits maladaptive loudness sensation leading to excessive signal contrast.

As in broadband noise the slope of the regression line is 0.1 dB/dB, the discovered effects corresponding to 0.3 and 0.4 dB in SRT for basilar-membrane compression and temporal fine-structure or temporal envelope resolution (DPOAE and ASSR, respectively) can be compensated only by a tenfold shift in threshold, corresponding then to 3-4 dB.

It thus might be tempting to hypothesize that a lower DPOAE acceptance rate (Fig. 9B) — explaining variance of OLSA-LP (Tab. 4) — is linked to good discrimination of phonemes with formant contrasts below PLL (/o/-/u/) in subjects with good PNOT (Fig.11, Tab. 6, Fig. 12). In other words, the higher contrast of the speech signal when there is no compression is advantageous for temporal fine-structure coding below the PLL limit and may occur on cost of envelope coding > PLL (Fig. 11, Tab. 6, Fig. 12).

A limitation to interpreting frequency-specific aspects of DPOAE I/O function acceptance rate in relation to phoneme discrimination is that the speech-intelligibility tests von BB, LP, and HP condition were presented at different intensities per frequency band because they were normalized so that the speech stimuli had the same overall perceptual levels. This may be a confounder especially with respect to recruitment.

### In conclusion

In conclusion, apart from the dominating threshold dependence, we discovered several effects which independently affect speech discrimination and differ depending on whether the speech signal is close to threshold (speech in quiet), or clearly supra-threshold (speech in noise). These effects were independent of age. As identified by comparison between DPOAE threshold and pure-tone threshold, neural firing rate adaptation at stimulus onset as well as cochlear synaptopathy contribute to speech comprehension in quiet. In noise, it appears that the recruitment phenomenon partially can counteract the discrimination deficits brought about by hearing loss due to reduced cochlear amplification. Differences in the nerve adaptation rate at stimulus onset in quiet and recruitment phenomenon in noise must therefore be re-included in the 50% differences in human speech understanding that were previously not explained by hearing thresholds and these should be considered as a new mechanistic principle behind the different coding principles predicted to dependent on PLL (Garrett et al., 2020; Moore, 2021).

## Acknowledgments and funding

This work was supported by the Deutsche Forschungsgemeinschaft DFG KN 316/13-1, DFG RU 713/6-1, KL 1093/12-1; ERA-NET NEURON JTC 2020: BMBF 01EW2102 CoSySpeech and FWO G0H6420N; IZKF Promotionskolleg of the Faculty of Medicine, University Hospital of Tübingen. We thank the audiometrists of the HNO-Klinik Tübingen for their expertise in recording audiograms.

## Author contributions

Designed research: M.K., L.R., S.W., M.M, E.D. E.G., S.V, D.B. Conducting experiments: J.S., K.D., M.R. Analyzed data: J.S., K.D., M.R., M.W., J.W., T.E., K.B, W.S., Wrote the paper: M.K., M.M., L.R., E.D., S.W. Edited the paper: M.K., L.R., S.W., M.M, E.D., E.G., D.B, S.V., C.B.

